# Programming of macrophages by apoptotic cancer cells inhibits cancer progression through exosomal PTEN and PPARγ ligands

**DOI:** 10.1101/217562

**Authors:** Yong-Bae Kim, Young-Ho Ahn, Jin-Hwa Lee, Jihee Lee Kang

## Abstract

Apoptotic cell clearance by phagocytes is essential in tissue homeostasis. We demonstrated that conditioned medium (CM) from macrophages exposed to apoptotic cancer cells inhibits epithelial-mesenchymal transition (EMT), migration, and invasion of cancer cells with the acquisition of cancer-stem–like traits. Apoptotic 344SQ (ApoSQ) cell-induced PPARγ activity in macrophages caused increased PTEN levels, secreted in exosomes. ApoSQ-exposed CM from PTEN knockdown cells failed to enhance PTEN in recipient 344SQ cells, restore cellular polarity, and exert anti-EMT and anti-invasive effects. The CM which deficient of PPARγ ligands could not reverse the suppression of PPARγ activity and PTEN and consequently failed to the prevent EMT process. Moreover, single injection of ApoSQ cells inhibited lung metastasis in syngeneic mice with enhanced PPARγ/PTEN signaling both in tumor-associated macrophages and tumor cells. PPARγ antagonist GW9662 reversed PTEN signaling and anti-metastatic effect. Thus, apoptotic cancer cell therapy may offer a new strategy for the prevention of metastasis.

## Introduction

Metastasis is a complex multistep process of cancer cell dissemination that is extremely difficult to treat. The outcome of cancer patients with metastatic disease has not improved in the past 30 years, in spite of the development of targeted therapies (Tevaarwerk et al., 2013). In one working hypothesis, metastasis is initiated by tumor cells that undergo epithelial-to-mesenchymal transition (EMT) in response to extracellular signals, leading to loss of polarity, detachment from neighboring cells, increased motility, invasion into surrounding matrix, and resistance to standard cytotoxic chemotherapy (Singh and Settleman, 2010).

*PTEN* (phosphatase and tensin homolog on chromosome ten), a powerful and multifaceted suppressor, is mutated in multiple types of cancer (Li et al., 1997) and has both phosphatase-dependent and -independent roles. *PTEN* encodes a dual-specificity phosphatase whose primary substrate is phosphatidylinositol 3,4,5 triphosphate (PIP3) (Cully et al., 2006). PTEN antagonizes phosphoinositide 3-kinase (PI3K) signaling and thereby affects several cellular processes, including growth, proliferation, and survival (Cantley, 2002, Wang et al., 2015). A number of clinical studies demonstrate that PTEN suppression or loss in advanced stage disease contributes to tumor invasion and metastasis (Wikman et al., 2012, Mulholland et al., 2012). PTEN knockdown in human colon cancer cells or prostate cancer cells leads to EMT induction, associated with invasion and metastasis (Bowen et al., 2009). In mice, *PTEN* loss results in neoplastic growth, in tumors and in the tumor microenvironment (Podsypanina et al., 1999, Trimboli et al., 2009). Two recent studies have found, surprisingly, that PTEN can be secreted unconventionally via exosome formation, and that the translational variant PTEN-long form can be secreted via an unknown mechanism from donor cells and enter neighboring cells (Hopkins et al., 2013, Putz et al., 2012). Like cytoplasmic PTEN, secreted PTEN has lipid phosphatase activity and antagonizes PI3K signaling in target cells, inducing tumor regression. Peroxisome proliferator-activated receptor gamma (PPARγ) is a potential PTEN transcription factor (Patel et al., 2001); its activation through ligands increases functional PTEN protein expression in various cancer cell lines, subsequently inhibiting Akt phosphorylation and cellular growth (Teresi et al., 2006, Zhang et al., 2006). Several *in vivo* studies have demonstrated that genetic alterations of PPARγ can promote tumor progression (Yu et al., 2010, Shen et al., 2012). These studies suggest the importance of PPARγ/PTEN signaling in cancer prevention.

Apoptotic cell clearance by tissue macrophages and non-professional phagocytes (efferocytosis) is essential in tissue homeostasis, immunity, and inflammation resolution. High levels of cell death can occur within the tumor environment, and clearance mechanisms for dying tumor cells can profoundly influence tumor-specific immunity. Previously, we demonstrated that *in vitro* and *in vivo* exposure of macrophages to apoptotic cells inhibits TGF-β1 or bleomycin-induced EMT in lung alveolar epithelial cells (Yoon et al., 2016). However, the effects of efferocytosis in the multistep process of cancer cell dissemination, leading to cancer metastasis, have not been studied thus far. Here, using *in vitro* 2D and 3D culture systems, we investigated whether the interaction of macrophages with dying cancer cells inhibits EMT in various epithelial cancer cells, and decreases cancer cell migration and invasiveness. We demonstrated that PTEN secretion in exosomes and the PPARγ ligands from macrophages exposed to apoptotic cancer cells block the multistep metastatic process. Furthermore, we provided *in vivo* evidence that subcutaneous injection of apoptotic cancer cells inhibits the numbers of visible lung metastases of the primary subcutaneous tumor via PPARγ/PTEN signaling.

## Results

### Interaction between macrophages and apoptotic cancer cells inhibits TGF-β1-induced EMT in cancer cells

To determine whether programming macrophages by apoptotic cancer cells inhibits EMT in epithelial cancer cells, 344SQ cells were treated with conditioned medium (CM) from RAW cells exposed to either apoptotic 344SQ murine lung adenocarcinoma cells (ApoSQ-exposed CM) or necrotic cells (NecSQ-exposed CM), along with TGF-β1. ApoSQ-exposed CM inhibited TGF-β1-induced EMT, based on morphological cellular alterations (Figure 1A), and EMT marker mRNA (Supplementary figure 1A) and protein (Figure 1B) expression profiles. In contrast, NecSQ-exposed CM did not exhibit anti-EMT effects. We confirmed the anti-EMT effects of various types of apoptotic cancer cell-exposed CM in the human non-small cell lung cancer (NSCLC) cell line A549 (Supplementary figure 1B-D) and other human cancer cells lines, including breast (MDA-MB-231), colon (COLO320HSR), and prostate (PC3) cancer cells (Supplementary figure 1E-G). In addition to RAW cells, CM from ApoSQ-exposed primary isolated mouse bone marrow-derived macrophages (BMDMs), or their IL-4-derived M2 phenotype, substantially inhibited TGF-β1-induced changes in EMT markers in 344SQ cells (Supplementary figure 1H,I). CM from blood monocyte-derived macrophages (MDMs) from healthy humans and NSCLC (adenocarcinoma) patients exposed to apoptotic A549 cells (ApoA) showed anti-EMT effects (Figure 1C; Supplementary figure 1J,K).

**Figure 1.**
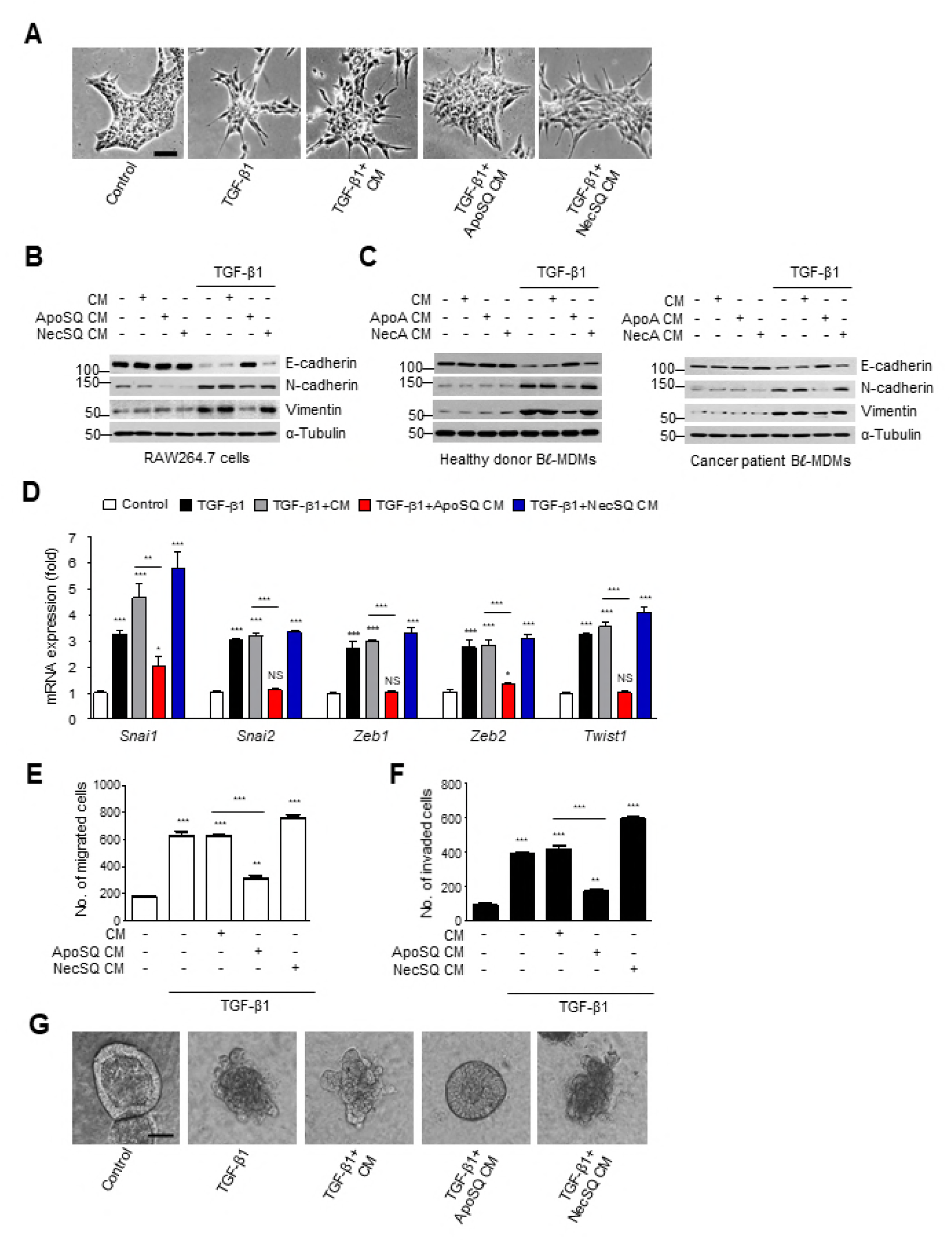
Interaction of macrophages and apoptotic cancer cells inhibits EMT, migration and invasion in cancer cells. (A, B and D–G) RAW cells were stimulated with apoptotic (ApoSQ) or necrotic (NecSQ) 344SQ cells for 24 h. (C) Blood monocyte-derived macrophages (MDMs) from healthy donor (*left*) or lung cancer patients (*right*) were stimulated with apoptotic (ApoA) or necrotic (NecA) A549 cells for 24 h. Conditioned medium (CM) was added to 344SQ cells (A, B, D–F) or A549 cells (C) with or without TGF-β1 (10 ng/ml) for 48 h. (A) Morphological changes in cells were examined by phase-contrast microscopy. (B and C) Immunoblot analysis of indicated proteins in whole-cell lysates. (D) Gene expression in 344SQ cell samples was measured by real-time PCR. The quantification of migrated (E) or invaded cells (F) in Boyden chambers. (G) 344SQ cells were cultured for 7 days in Matrigel before exposure to CM with TGF-β1 for 72 h and photographed under phase-contrast microcopy. \**P* < 0.05, \*\**P* < 0.01 and \*\*\**P* < 0.001. Data are from one experiment representative of three independent experiments (A, B, G) or three donors (C) with similar results, or from three independent experiments (mean ± s.e.m.; D–F). Scale bars: 100 µm (A and G).

However, neither CM from nor direct exposure to epithelial cancer cells inhibited TGF-β1-induced EMT marker changes (Supplementary figure 2A-D), indicating that cancer cell EMT inhibition requires bioactive mediators secreted by professional phagocytes, such as macrophages, which are functionally altered by apoptotic cancer cell stimulation.

TGF-β-induced EMT is achieved through the well-orchestrated actions of the Snai, ZEB, and Basic helix-loop-helix transcription factor families (Xu et al., 2009). We observed that ApoSQ or ApoA-exposed CM markedly inhibited TGF-β1-induced *Snai1/2, Zeb1/2*, and *Twist1* mRNA expression (Figure 1D; Supplementary figure 3A), whereas control, NecSQ-, or NecA-exposed CM did not. Intracellular signaling studies (Supplementary figure 3B) show that Smad-dependent TGF-β signaling and the ERK pathway were not affected (Supplementary figure 3C-E). However, ApoSQ-exposed CM partially blocked Smad-independent TGF-β signaling, including the p38 MAP kinase and Akt pathways, in 344SQ cells (Supplementary figure 3F,G).

### Interaction of macrophages with apoptotic cancer cells inhibits TGF-β1-induced cancer cell migration and invasion

Acquisition of mesenchymal state by malignant cancer cells is associated with decreased cell-cell adhesion, and increased migratory and invasive properties, which are crucial for metastasis (Lamouille et al., 2014). These processes are consistent with the acquisition of a cancer stem-like cell phenotype also known as ‘stemness’ or cancer stem cell characteristics (Brabletz et al., 2005). Our data show that ApoSQ-or ApoA-exposed CM prevented TGF-β1-induced cancer cell migration and invasion (Figure 1E,F; Supplementary figure 3H,I), whereas control, NecSQ-, or NecA-exposed CM did not. In addition, TGF-β1-induced gene-level enhancement of cancer stem-like cell markers, such as *CD90, Oct4, CD44, CD133,* or *ALDH1A,* was reduced by treatment with ApoSQ or ApoA-exposed CM (Supplementary figure 3J,K). Moreover, 3D Matrigel culture confirmed the anti-invasive effect of ApoSQ-exposed CM, which caused 344SQ cells to recover their lost polarity and acinus-like colonies, and caused invasive projection by TGF-β1 (Figure 1G).

### PPARγ-dependent PTEN secretion in exosomes from macrophages exposed to apoptotic cancer cells

PPARγ expression and activity were enhanced in macrophages exposed to apoptotic cells *in vitro* and *in vivo* (Freire-de-Lima et al., 2006, Yoon et al., 2015). We hypothesized that PPARγ-dependent PTEN production in macrophages in response to apoptotic cancer cells plays a crucial role in the anti-EMT effect in cancer cells. To prove this, we examined PPARγ expression and activity, and PTEN production, in ApoSQ-exposed RAW cells. PPARγ mRNA and protein expression, and its activity, were markedly enhanced (Figure 2A; Supplementary figure 4A-D). *PPARγ* mRNA expression was enhanced in an apoptotic cell number-dependent manner (Supplementary figure 4A). In parallel, PTEN mRNA and protein levels were markedly increased (Figure 2B; Supplementary figure 4B,C). *PTEN* mRNA expression in blood MDMs from healthy humans or lung cancer patients was also enhanced by ApoA exposure (Figure 2C). NecSQ or NecA did not exert these effects (Figure 2C; Supplementary figure 4A-C). PPARγ inhibition by GW9662 (Supplementary figure 4D-F) or PPARγ knockdown with siRNA (Figure 2D) reversed the PTEN mRNA and protein abundance in ApoSQ-exposed RAW cells. Similar PPARγ-dependent PTEN induction was shown in naive BMDMs and M2-phenotype BMDMs exposed to ApoSQ (Supplementary figure 4G,H). These data suggest that this important protein is transcriptionally upregulated by enhanced PPARγ expression and activation in macrophages, upon stimulation with apoptotic cancer cells.

**Figure 2.**
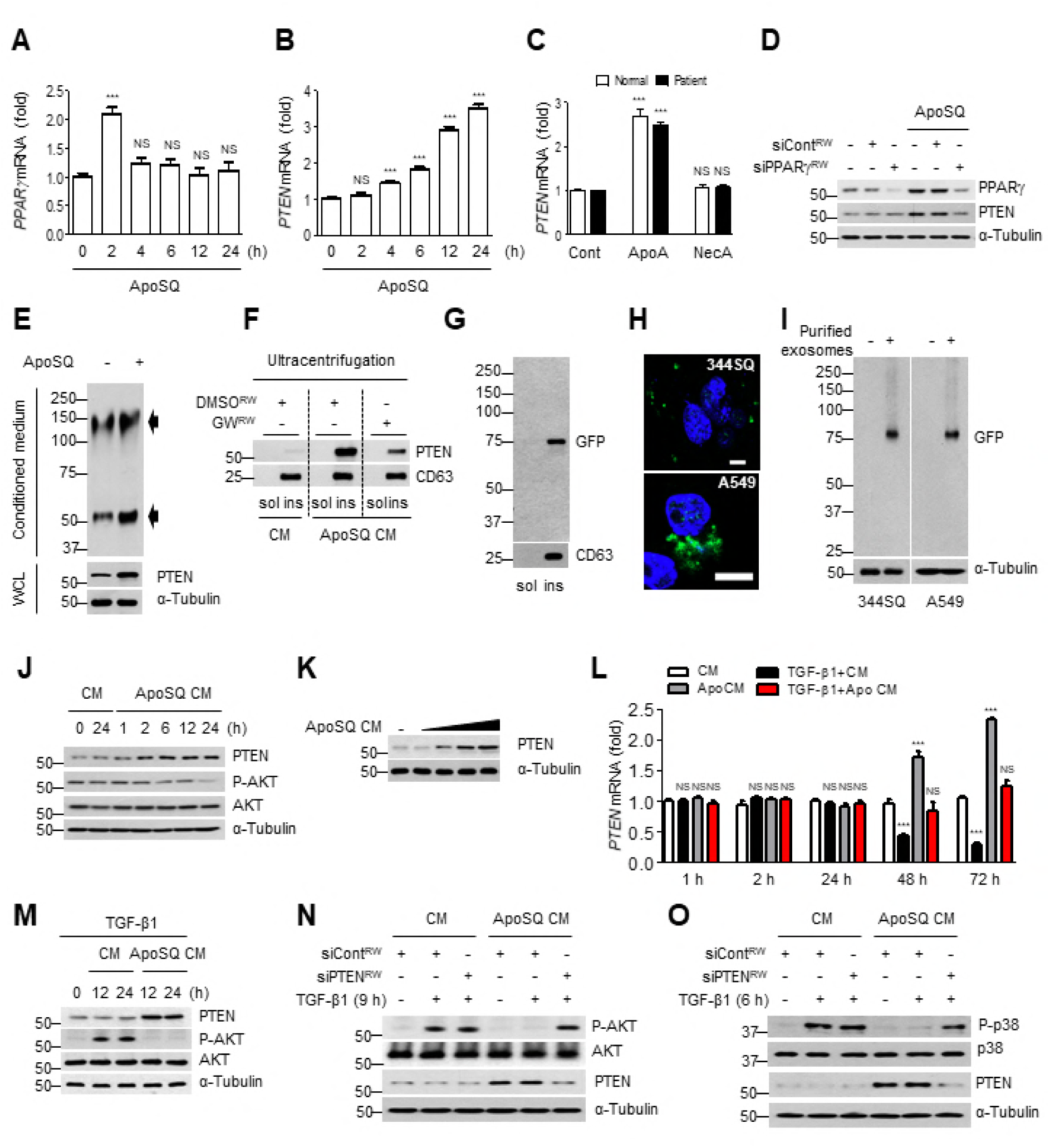
PPARγ-dependent PTEN secretion in exosomes alters signaling in recipient cells. qPCR analysis of *PPARγ* and *PTEN* mRNAs in RAW cells exposed to apoptotic 344SQ cells (ApoSQ) for the indicated times (A and B) or *PTEN* mRNA in blood MDMs from healthy donors or lung cancer patients exposed to apoptotic (ApoA) or necrotic (NecA) A549 cells for 24 h (C). (D) Immunoblot analysis of indicated proteins in RAW cells transfected with siRNA of PPARγ before ApoSQ stimulation for 24 h. (E) Immunoblot analysis of PTEN with whole-cell lysates from mouse BMDMs stimulated with ApoSQ (WCL) and for secretion levels of PTEN using affinity pull-down in conditioned medium (CM). PTEN immunoprecipitates were separated by SDS-PAGE in non-reducing conditions. Arrows indicate immunoglobulin heavy/light chain complex (up) and PTEN (down). (F) CM from RAW cells pretreated with 10 µM GW9662 before ApoSQ stimulation for 24 h was fractioned by ultracentrifugation, and the soluble (sol) and insoluble (ins) fractions were immunoblotted with anti-PTEN or CD63 antibodies. (G) Immunoblot analysis of GFP-PTEN and CD63 in the sol and ins fractions after ultracentrifugation of CM from human macrophage cell line (hMϕ) overexpressing GFP-PTEN exposed to ApoA. (H) Direct fluorescence of 344SQ and A549 cells 24 h after treatment with harvested exosomes containing GFP-PTEN using confocal microcopy. Scale bars: 20 µm (I) Lysates from 344SQ and A549 cells after incubation with harvested exosomes from hMϕ overexpressing GFP-PTEN were subjected to western blotting analysis. (J) Immunoblot analysis of indicated proteins in 344SQ cell lysates after incubation with CM for the indicated times. (K) Immunoblot analysis of PTEN in 344SQ cells after incubation with ApoSQ-exposed CM at various dilutions (from 1/5 to 1)with control CM for 12 h. qPCR analysis of *PTEN* mRNA (L) and Immunoblot analysis of indicated proteins (M) in 344SQ cells after addition of CM. (N and O) Immunoblot analysis of indicated proteins in 344SQ cells after addition CM from RAW cells transfected with PTEN siRNA, in the presence of TGF-β1. NS, not significant; \*\*\**P* < 0.001. Data are from three independent experiments (mean ± s.e.m. in A, B and L), three donors (mean ± s.e.m. in C), or one experiment representative of three independent experiments with similar results (D-K, M-O).

Interestingly, GW9662 treatment in RAW cells significantly reversed the anti-EMT effects of ApoSQ-exposed CM (Supplementary figure 4I-L). PPARγ-dependent PTEN production in this experimental context might be a candidate for the acquisition of these anti-EMT and anti-invasive effects of the CM, if PTEN can be secreted in exosomes and secreted PTEN internalized by recipient cells, with consequent functional activity. To confirm this assumption, we first examined whether the enhanced PTEN protein levels could be secreted in exosomes from macrophages. Transmission electron microscopy revealed microvesicles in RAW cells treated with ApoSQ (Supplementary figure 4M). In addition to ApoSQ-induced PTEN expression in BMDM whole-cell lysates, we observed strong PTEN expression in ApoSQ-exposed CM (Figure 2E). Moreover, exosomes were isolated by sequential ultracentrifugation (Putz et al., 2012) and used for western blotting analysis using anti-PTEN and exsosomal marker CD63 antibodies. Importantly, PTEN was recovered in the insoluble fraction with the exosomal marker CD63, confirming the presence of PTEN in ApoSQ-exposed CM, whereas GW9662 treatment reduced PTEN abundance (Figure 2F).

### PTEN alters signaling, promotes cellular polarity, and inhibits EMT upon entry into 344SQ cells

To ascertain entry of PTEN-bearing exosomes secreted from macrophages to the recipient cancer cells, CM from human macrophage cells overexpressing GFP-PTEN exposed to ApoA was subjected to sequential ultracentrifugation for the harvesting of exosomes. We confirmed the presence of exosomal PTEN in the insoluble fraction through western blotting analysis using anti-GFP and CD63 antibodies (Figure 2G). After treating cells, such as 344SQ and A549 cells, with the harvested exosomes for 24 h, GFP fluorescence was detected in the cells using confocal microscopy (Figure 2H), indicating uptake of exosomal GFP-PTEN. This observation was confirmed using western blotting with anti-GFP antibodies (Figure 2I).

PTEN protein levels in recipient 344SQ cells increased immediately and remained so up to 24 h, and basal Akt phosphorylation decreased reciprocally over 6 h, following treatment with ApoSQ-exposed CM (Figure 2J). Moreover, PTEN abundance was inversely proportional to the dilution of ApoSQ-exposed CM (Figure 2K), although its mRNA abundance was not affected until 24 h after ApoSQ-exposed CM treatment in the absence or presence of TGF-β1 (Figure 2L). TGF-β1 itself did not affect basal PTEN protein abundance for 24 h, and thereafter caused it to decrease in 344SQ cells (Supplementary figure 4N). Reciprocal to PTEN abundance, TGF-β1-induced Akt phosphorylation decreased until 24 h after ApoSQ-exposed CM treatment (Figure 2M). However, PTEN enhancement, and the effective reduction of Akt phosphorylation or p38 MAP kinase, were not seen with ApoSQ-exposed CM from PTEN knockdown RAW cells (Supplementary figure 4O; Figure 2N,O). These data suggest that entered PTEN is functionally active to alter basal Akt signaling and TGF-β1-induced non-Smad signaling.

We next investigated whether PTEN is recruited to the plasma membrane after treatment with ApoSQ-exposed CM to sustain cell polarity. Twelve hours after ApoSQ-exposed CM treatment, PTEN protein was enhanced in the plasma membrane-enriched fraction in 344SQ cells (Figure 3A). Pretreatment with the PTEN-selective inhibitor SF1670, which elevates intracellular PIP3 signaling (Li et al., 2014), reversed the inhibition of basal Akt phosphorylation in the plasma membrane-enriched fraction. These data suggest that PTEN is recruited to the plasma membrane and functionally inhibits Akt activation.

**Figure 3.**
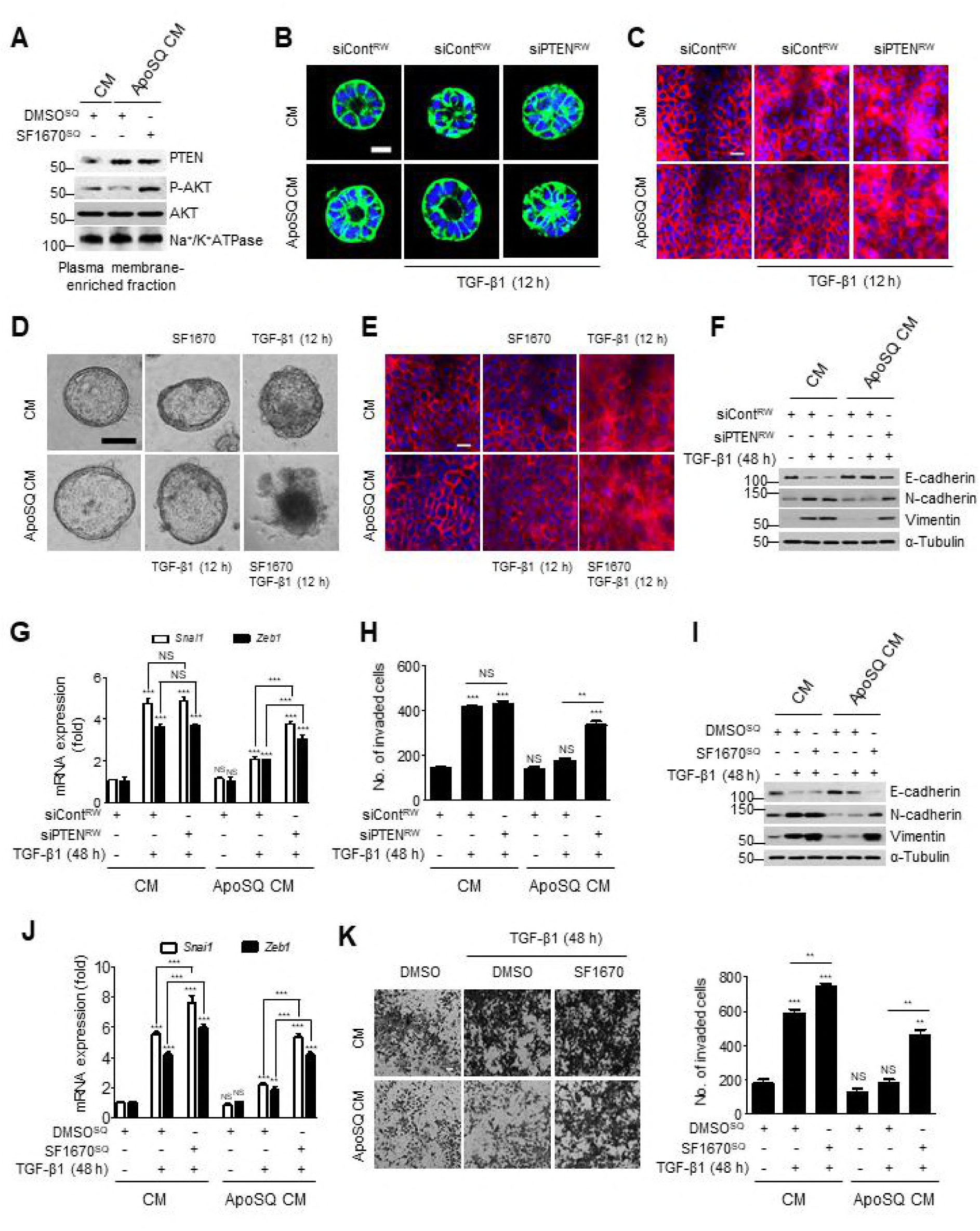
Role of internalized PTEN in retaining polarity, and inhibition of EMT and invasion of 344SQ cells. (A) 344SQ cells were pretreated with SF1670 (10 μM) for 1 h and then treated with CM or ApoSQ CM for 12 h. Immunoblot analysis of indicated proteins in plasma membrane-enriched fractions of the cells. (B, C and F-H) RAW cells were transfected with siRNA of PTEN before ApoSQ cell stimulation for 24 h. (B) Normal acini of 344SQ cells were grown in 3D Matrigel containing CM and stained with anti-γ-catenin (green) and DAPI. CM was added to 344SQ cells with or without TGF-β1 for 12 h (C) and for 48 h (F-H). 344SQ cells were pretreated with SF1670 (10 μM) for 1 h and then treated with CM with or without TGF-β1 for 12 h (D and E) and 48 h (I-K). (C and E) Immunofluorescence staining for β-catenin (red) and nuclei (DAPI) in 344SQ cells. (D) Phase-contrast images of cells grown in 3D Matrigel cultures in the indicated conditions. (F and I) Immunoblot analysis of EMT markers in cell lysates. (G and J) qPCR analysis of *Snai1* and *Zeb1* mRNAs in 344SQ cells. The invaded cell numbers (H and K *right*) and the cells visualized by phase-contrast microscopy to analyze their invasive ability using Matrigel-coated Transwell (K *left*). NS: not significant; \*\**P* < 0.01 and \*\*\**P* < 0.001. Data are from one experiment representative of three independent experiments with similar results (A-F, I and K *left*), from three independent experiments (mean ± s.e.m., in G and J), or from three fields from replicate wells (H and K *right*). Scale bars: 20 μm (B, C and E) and 100 μm (D and K).

Loss of PTEN function prevents normal apical surface and lumen development in 3D-culture (Martin-Belmonte et al., 2007). Thus, to evaluate the role of PTEN in cell polarity, 344SQ cells grown in 3D Matrigel were treated with CM from PTEN knockdown RAW cells exposed to ApoSQ, stained with anti-β-catenin (green), and examined by confocal microscopy. Treatment with ApoSQ-exposed CM from control siRNA-transfected macrophages prevented TGF-β1-induced interference, with the formation of polarized acinar structures by 344SQ cells during relatively early exposure (12 h after TGF-β1 treatment) (Figure 3B). ApoSQ-exposed CM from PTEN knockdown macrophages did not exert this inhibitory effect. Moreover, TGF-β1-induced disruption of cell-to-cell contacts was prevented by ApoSQ-exposed CM treatment, whereas CM from PTEN knockdown macrophages suppressed this preventive effect (Figure 3C). Similarly, pretreatment of 344SQ cells with SF1670 inhibited ApoSQ-exposed CM-induced restoration of cellular polarity in 3D Matrigel culture (Figure 3D) and cell-to-cell contact, as evidenced by confocal microscopy after anti-β-catenin staining (red) (Figure 3E).

Next, we examined whether PTEN contributes to the late-phase anti-EMT and anti-invasive effects of ApoSQ-exposed CM. PTEN knockdown in RAW cells substantially inhibited ApoSQ-exposed CM-induced EMT marker changes (Figure 3F,G), and invading cell number reduction (Figure 3H), 48 h after TGF-β1 treatment. Similarly, PTEN inhibition by SF1670 in 344SQ cells reversed the anti-EMT and anti-invasion effects of ApoSQ-exposed CM (Supplementary figure 3P; Figure 3I-K). These data indicate that PTEN or its signaling mediates the anti-EMT and anti-invasive effects of ApoSQ-exposed CM in the late phase in TGF-β1-stimulated 344SQ cells.

### Secreted PPARγ ligands from macrophages mediate anti-EMT effects in 344SQ cells via PPARγ/PTEN signaling

Apoptotic cells stimulate the production of the identified PPARγ ligands, such as 15-lipoxygenase-dependent 15-hydroxyeicosatetraenoic acid (HETE), lipoxin A4, and PGD_2_ synthase-dependent 15-deoxy-Δ12,14- prostaglandin J_2_ (15d-PGJ_2_) (Fujimori et al., 2012, Korns et al., 2011). Consistent with these findings, we observed enhanced production of these PPARγ ligands in ApoSQ-exposed CM (Figure 4A-C). However, viable or necrotic 344SQ cells did not show these effects. PPARγ activity gradually increased in 344SQ cells over 36–72 h after treatment with ApoSQ-exposed CM in the absence of TGF-β1 (Figure 4D). Consistent with previous findings (Wei et al., 2010), *PPARγ* mRNA abundance (Supplementary figure 5A,B) and activity (Figure 4E,F) were suppressed upon TGF-β1 stimulation, as were PTEN mRNA (Figure 2L; Supplementary figure 5C,D) and protein levels at 48 h (Figure 4G,H). However, ApoSQ-exposed CM treatment reversed these reductions. To confirm that the PPARγ activity increase is mediated by secreted PPARγ ligands, RAW cells were pretreated with the 15-lipoxygenase inhibitor PD146176, or transfected with siRNA of lipocalin-type PGD synthase (L-PGDS), before ApoSQ treatment. Under these conditions or with L-PGDS protein knockdown (Figure 4I), 15-HETE, lipoxin A4, PGD_2_, and 15d-PGJ_2_ production was substantially reduced (Supplementary figure 5E-H), whereas PPARγ activity was not in ApoSQ-exposed RAW cells (Supplementary figure 5I,J), suggesting no autocrine effect of these ligands on PPARγ in RAW cells. Notably, treatment with this CM, deficient of PPARγ ligands, could not effectively reverse the TGF-β1-induced suppression of *PPARγ* mRNA (Supplementary figure 5A,B) and activity (Figure 4E,F), as well as PTEN mRNA (Supplementary figure 5C,D) and protein levels (Figure 4G,H), 48 h after TGF-β1 treatment. Consequently, late EMT processes were not effectively prevented (Figure 4J,K; Supplementary figure 4K,L). These data suggest that macrophage secretion of these ligands mediates anti-EMT effects through enhanced PPARγ/PTEN signaling in recipient 344SQ cells.

**Figure 4.**
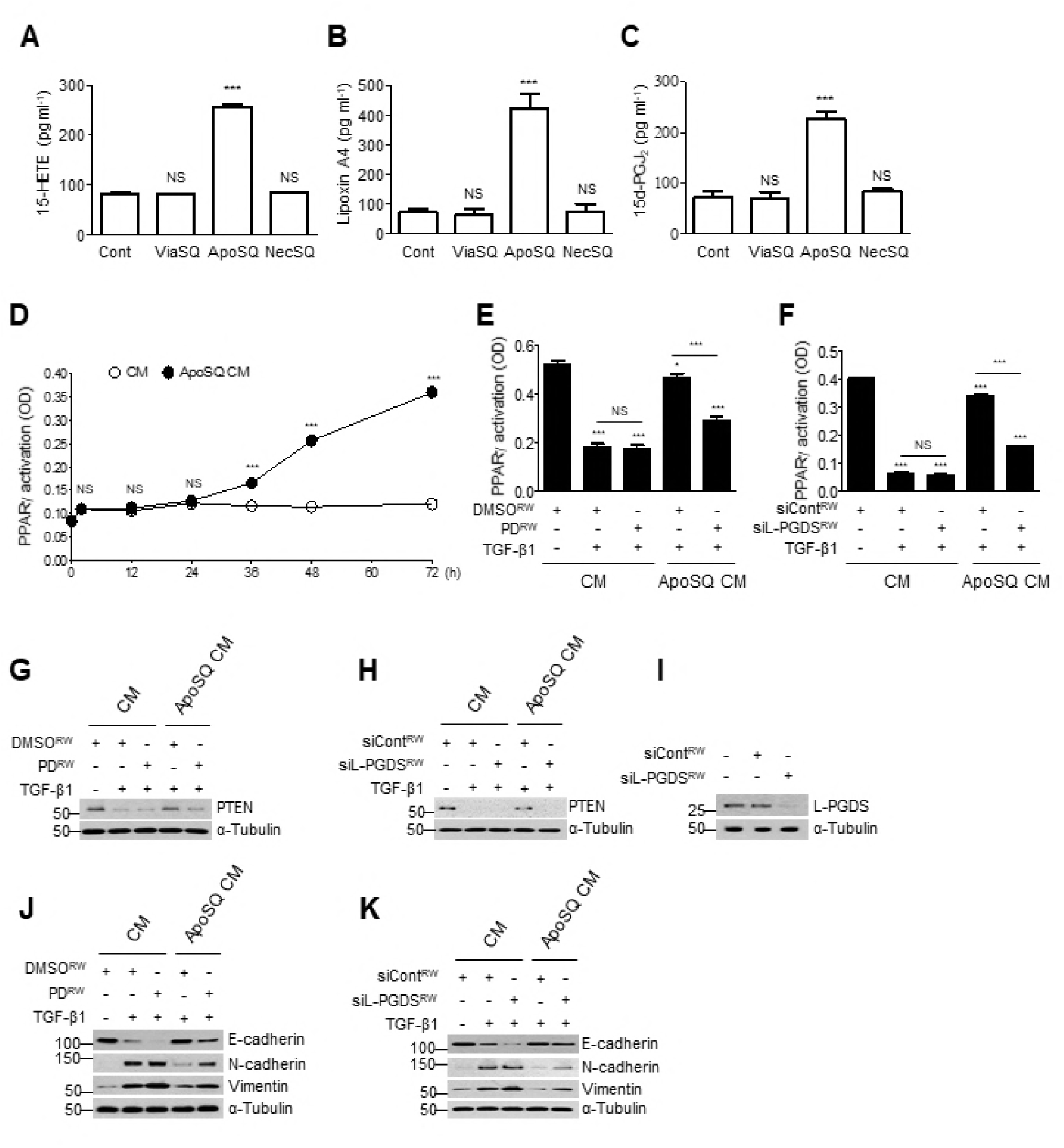
Macrophage secretion of 15-HETE, lipoxin A4, and 15d-PGJ_2_ mediates anti-EMT effects. (A-C) ELISA of 15-HETE, lipoxin A4 and 15d-PGJ_2_ in conditioned medium (CM) from RAW cells alone, RAW/viable 344SQ cell (ViaSQ), RAW/apoptotic 344SQ cell (ApoSQ), or RAW/necrotic 344SQ cell (NecSQ) co-cultures. (D) Time course of PPARγ activation in 344SQ cells with CM or ApoSQ CM at the indicated times. RAW cells were pretreated with 10 μM PD146176 (PD) for 1 h (E, G and J) or transfected with siRNA against lipocalin-type prostaglandin D synthase (siL-PGDS) for 48 h (F, H, I and K) before stimulation with ApoSQ cells for 24 h. (E–H, J and K) CM was added to 344SQ cells with TGF-β1 (10 ng/ml) for 48 h. PPARγ activation (E and F), immunoblot analysis of indicated protein expression in 344SQ cells (G, H, J and K) and L-PGDS expression in RAW cells (I). NS: not significant; \**P* < 0.05, \*\**P* < 0.01 and \*\*\**P* < 0.001. Data are from three independent experiments (mean ± s.e.m. in A–F), or from one experiment representative of three independent experiments with similar results (G–K).

### Exogenous treatment of cancer cells with ligands inhibits EMT via enhanced PPARγ/PTEN signaling

To confirm that 15-HETE, lipoxin A4, and 15d-PGJ_2_ act in a paracrine manner to induce anti-EMT effects through enhanced PPARγ/PTEN signaling, we investigated the effects of these soluble mediators on 344SQ cells at basal (80, 73, and 73 pg/ml, respectively) and stimulatory (258, 422, and 226 pg/ml, respectively) concentrations. Stimulatory concentrations of all these ligands combined enhanced PPARγ activity after 36 h, whereas basal concentrations exerted no effect (Supplementary figure 6A). As expected, each ligand partially inhibited late-phase TGF-β1-induced EMT process at its stimulatory, but not basal, concentration (Supplementary figure 6B,C). In parallel, TGF-β1-induced reduction of *PPARγ* mRNA abundance and activity (Supplementary figure 6D,E), and PTEN mRNA and protein abundances in 344SQ cells (Supplementary figure 6F,G), were reversed at stimulatory, but not basal, concentrations. In contrast, TGF-β1-induced non-Smad signaling, such as p38 MAP kinase and Akt phosphorylation, was not affected (Supplementary figure 6H,I)

### PTEN sources in recipient 344SQ cells include internalized PTEN and ligand-dependent PPARγ activity

We further investigated sources of PTEN signaling in 344SQ cells over time following ApoSQ-exposed CM treatment, using pharmacological strategies. The effect of ApoSQ-exposed CM on PTEN mRNA abundance at 48 h in the absence (Supplementary figure 7A) or presence (Supplementary figure 7B) of TGF-β stimulation was inhibited by GW9662 treatment in 344SQ cells, but not in RAW cells, indicating PPARγ-dependent PTEN induction in 344SQ cells. However, enhanced PTEN protein abundance in 344SQ cells in the presence of TGF-β1 at 12 h after ApoSQ-exposed CM treatment was reduced by GW9662 treatment in RAW cells, but not in recipient 344SQ cells, indicating that the PTEN is originated from PPARγ-dependent PTEN induction in RAW cells (Supplementary figure 7C). Associated with PTEN reduction in 344SQ cells, GW9662 treatment in RAW cells, but not in 344SQ cells, could inhibit cell polarity restoration by ApoSQ-exposed CM at 12 h (Supplementary figure 7D). However, ApoSQ-exposed CM from RAW cells treated with PD146176 or transfected with L-PGDS-siRNA did not alter PTEN protein abundance in 344SQ cells at 12 h after TGF-β1 stimulation (Supplementary figure 7E,G) and accordingly, had no effect on retaining cell polarity (Supplementary figure 7F,H). These data indicate that PTEN sources in recipient 344SQ cells include internalized PTEN in the early phase, and ligand-dependent PPARγ activity in the late phase, after ApoSQ-exposed CM treatment.

### Apoptotic cell treatment suppresses metastasis and enhances PPARγ/PTEN signaling *in vivo*

To explore the effects of apoptotic cancer cells in mouse metastasis models, we injected syngeneic (129/Sν) immunocompetent mice subcutaneously with highly metastatic 344SQ cells and allowed them to grow for 6 weeks (Figure 5A *left*). Subcutaneous ApoSQ administration two days after 344SQ cell injection did not significantly alter primary tumor size (Figure 5A *right*,B), but diminished tumor nodule number in lungs and the number of mice with visible lung metastasis 6 weeks after 344SQ cell injection (Figure 5C-E). ApoSQ injection increased the levels of *PPARγ* and its target molecules *PTEN* and *CD36*, but reduced *Snai1 and Zeb1* mRNA levels (Figure 5F-J), as well as Akt phosphorylation (Figure 5K).

**Figure 5.**
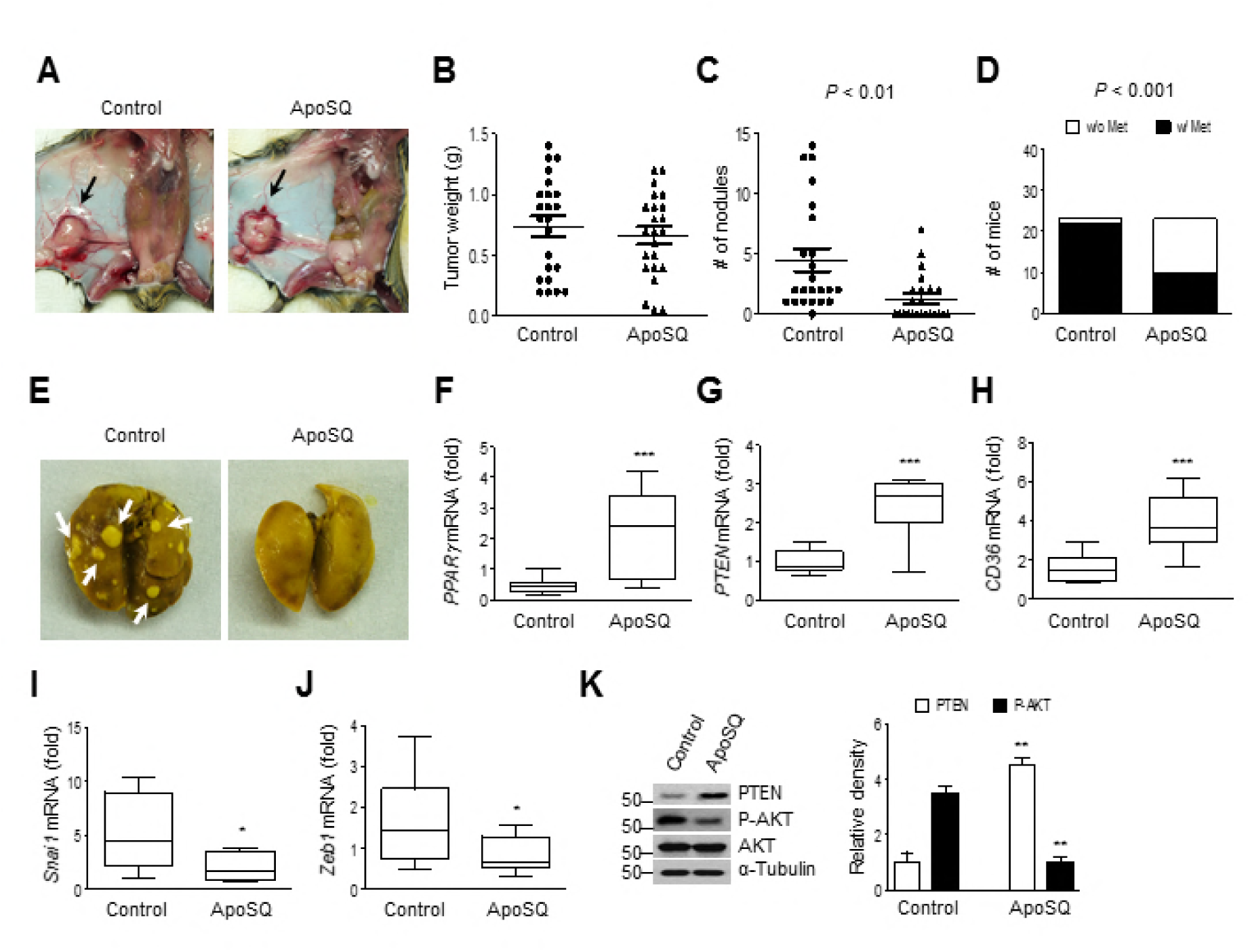
Anti-metastatic effects of apoptotic cancer cell injection in mice. Apoptotic 344SQ cells (ApoSQ) were subcutaneously injected in the skin lesion 2 days after subcutaneous injection of 344SQ cells into syngeneic (129/Sν) mice (n=23 per group). Mice were necropsied 6 weeks later. (A) Representative images of primary tumors (black arrows). Scatter plots of primary tumor weight (B) and numbers of metastatic pulmonary nodules (C). *P* values were determined by Student’s *t* test. (D) Bar graph indicates numbers of mice with (w/) or without (w/o) visibly determined metastases (Met). Metastasis incidence *P* value (Fisher’s exact test). (E) Representative images of metastatic (left) or non-metastatic lungs (right). White arrows indicate metastatic pulmonary nodules. qPCR analysis (F–J) and immunoblot analysis of indicated protein expression (K) in primary tumors. \**P* < 0.05, \*\**P* < 0.01 and \*\*\**P* < 0.001 (Student’s *t* test). The box represents the 25th to 75th percentile and whisker plots represent the minimum and the maximum percentiles. Data are from mice with lung metastasis [control; n=14 (F–J) or n=5 (K)] and without lung metastasis [ApoSQ; n=8 (F–J) or n=5 (K)] (mean ± s.e.m. in F–K).

Moreover, we found enhanced mRNA expression of *PPARγ* and its target genes, including *PTEN and CD36*, in tumor-associated macrophages (TAM) isolated from primary tumor after ApoSQ injection, compared to the control (Figure 6A). Using confocal microscopy, enhanced protein expression of these molecules (red) was confirmed in TAM stained with F4/80 (green) (Figure 6B-D). Similar to what we observed in culture in vitro, secretion of PPARγ ligands, such as 15-HETE, lipoxin A4 and 15d-PGJ_2_, was enhanced in culture media of TAM (Figure 6E). In parallel, mRNA and protein abundances of PPARγ and PTEN were also increased in remainder cells lacking TAM, mostly tumor cells, following ApoSQ injection (Supplementary figure 8A-D).

**Figure 6.**
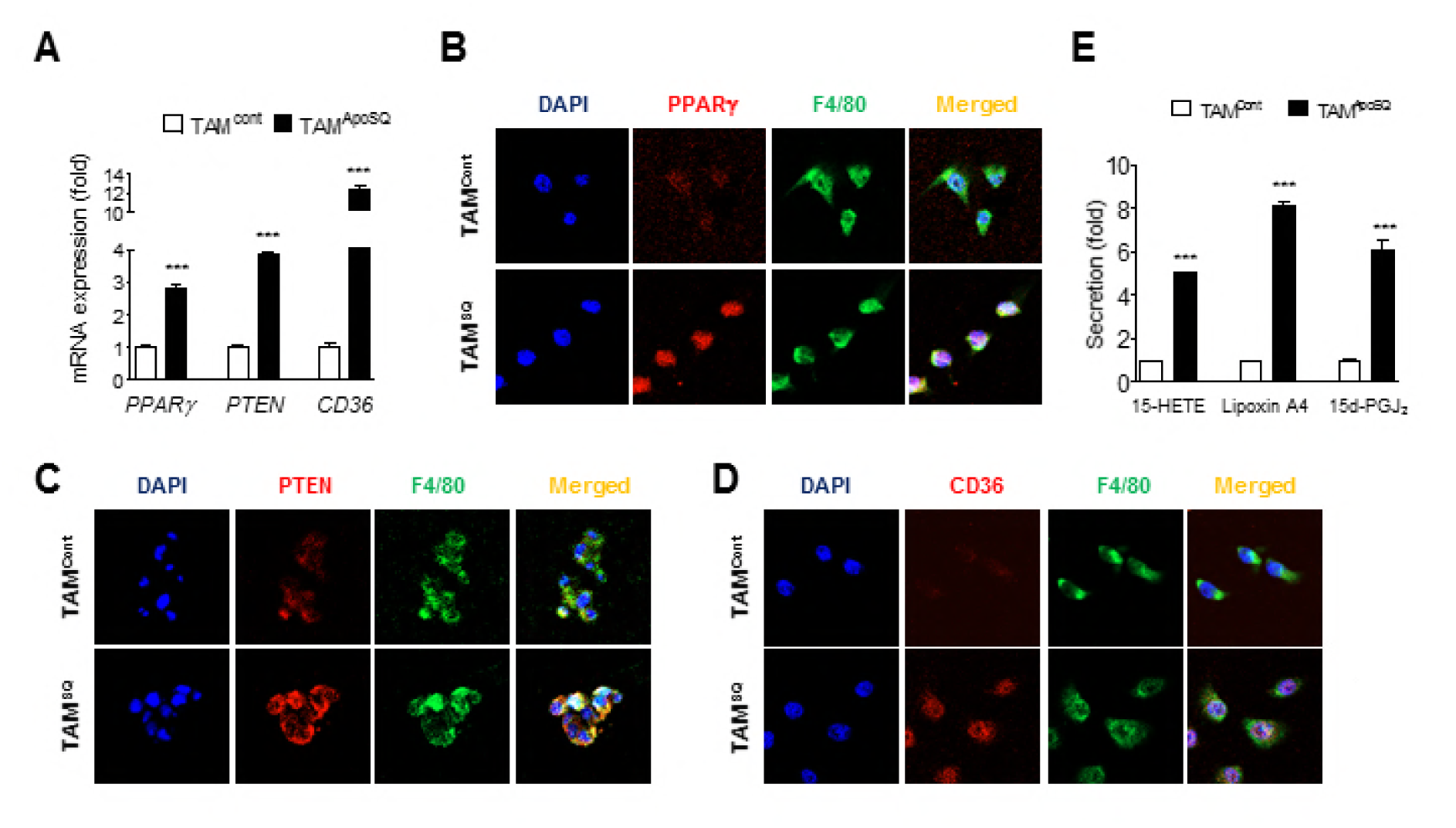
Expression of PPARγ and PTEN and secretion of PPARγ ligands in isolated TAM. (A) qPCR analysis of TAM isolated primary tumors (n=8). (B-D) Representative confocal images of TAM stained with anti-PPARγ (red), anti-PTEN (red), anti-CD36 (red) and anti-F4/80 (green). (E) ELISA of 15-HETE, lipoxin A4 and 15d-PGJ_2_ in TAM culture (n=8). \*\*\**P* < 0.001 (Student’s *t* test). Data are from mice with lung metastasis [control; n=8 (A–E)] and without lung metastasis [ApoSQ; n=8 (A–E)] (mean ± s.e.m. in A–E).

Immunohistochemistry of serial sections of primary tumor tissue confirmed enhanced PPARγ (*green*, Figure 7A,B), PTEN (*green*, Figure 7E,F), and CD36 expression (*red*, Figure 7H,I) upon ApoSQ injection. In particular, PPARγ (Figure 7D), PTEN (Figure 7G), and CD36 (Figure 7J) expression in F4/80-positive cells (*red*) was also markedly enhanced by apoptotic cell injection, which reflects apparent PPARγ, PTEN, and CD36 induction in tumor-infiltrating macrophages, as the F4/80-positive macrophage intensity was not different (Figure 7C).

**Figure 7.**
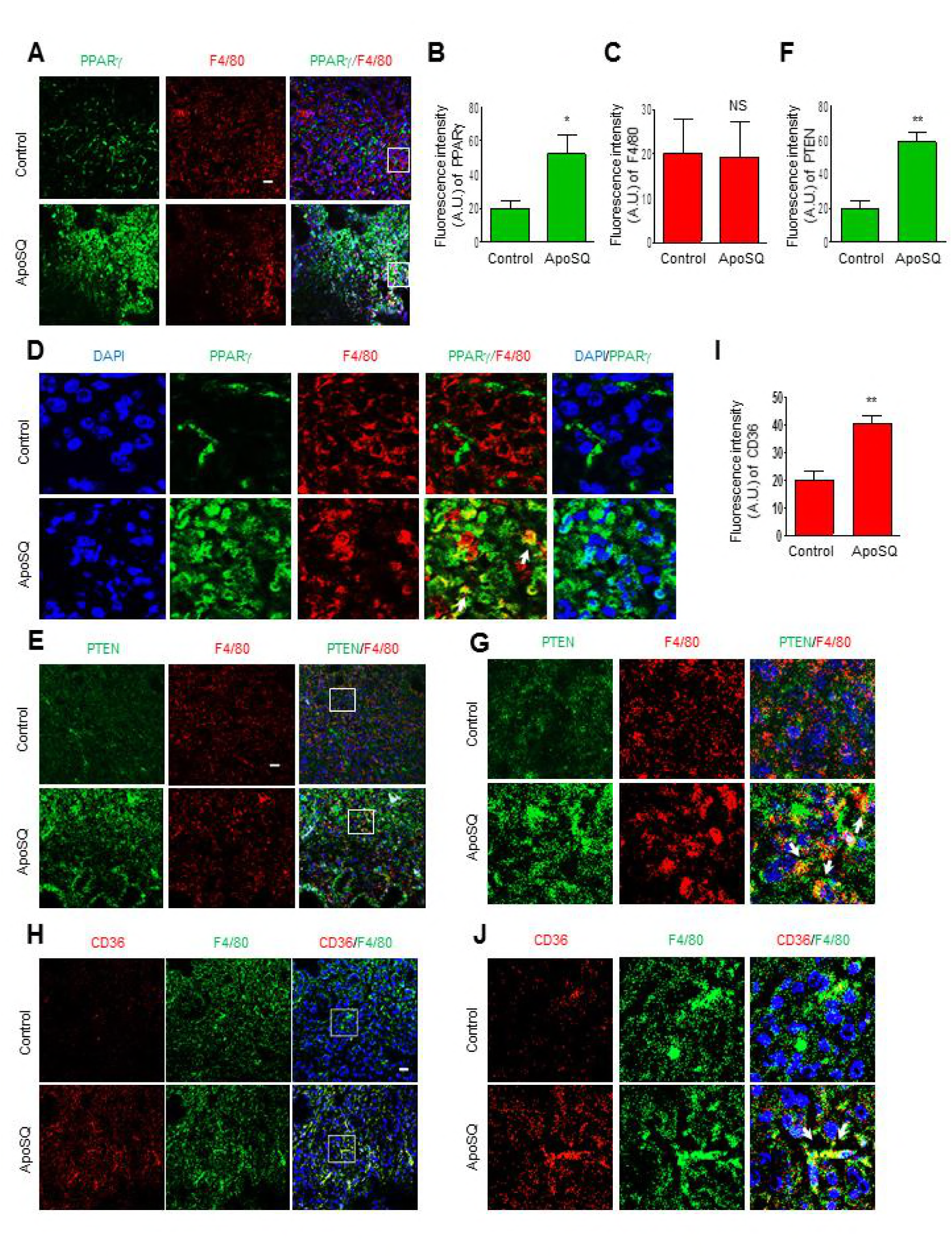
PPARγ and PTEN expression in primary tumors and tumor-infiltrating macrophages. Apoptotic 344SQ cells (ApoSQ) were subcutaneously injected into the skin lesion 2 days after subcutaneous injection of 344SQ cells into syngeneic (129/Sν) mice. Mice were necropsied 6 weeks later. Representative confocal images of primary tumors stained with anti-PPARγ (green) and anti-F4/80 (red) (A and D), anti-PTEN (green) and anti-F4/80 (red) (E and G), or anti-CD36 (red) and anti-F4/80 (green) (H and J), and the DNA-binding dye DAPI. (B, C, F and I) Measurements of fluorescence intensity on full-size images. (D, G and J) ROIs from white squares on the low magnification images of (A, E and H), respectively. Arrows indicate the localization of PPARγ, PTEN, and CD36 in macrophages. NS: not significant, \**P* < 0.05 and \*\**P* < 0.01. Data are representative images from five mice per group (A, E and H) or from independent experiments with five mice per group (mean ± s.e.m. in B, C, F and I). Scale bars: 100 μm (A, E and H).

To investigate the early response of macrophages to ApoSQ injection in wild-type mice injected subcutaneously with 344SQ cells, with respect to PPARγ and PTEN induction, double immunofluorescence staining with PPARγ, PTEN, or F4/80 Ab was performed on cryosections derived from the skin lesion. PPARγ (Supplementary figure 9A) and PTEN (Supplementary figure 9B) staining were markedly enhanced in F4/80-positive macrophages at 24 h following subcutaneous ApoSQ injection, compared to the control group.

Of note, in the ApoSQ injection group, we observed PTEN-bearing exosomes in the tumor sections using confocal microscopy (Figure 8A). In addition, increased PTEN expression was detected in the harvested exosomes from serum and ascitic fluid (Figure 8B,C).

**Figure 8.**
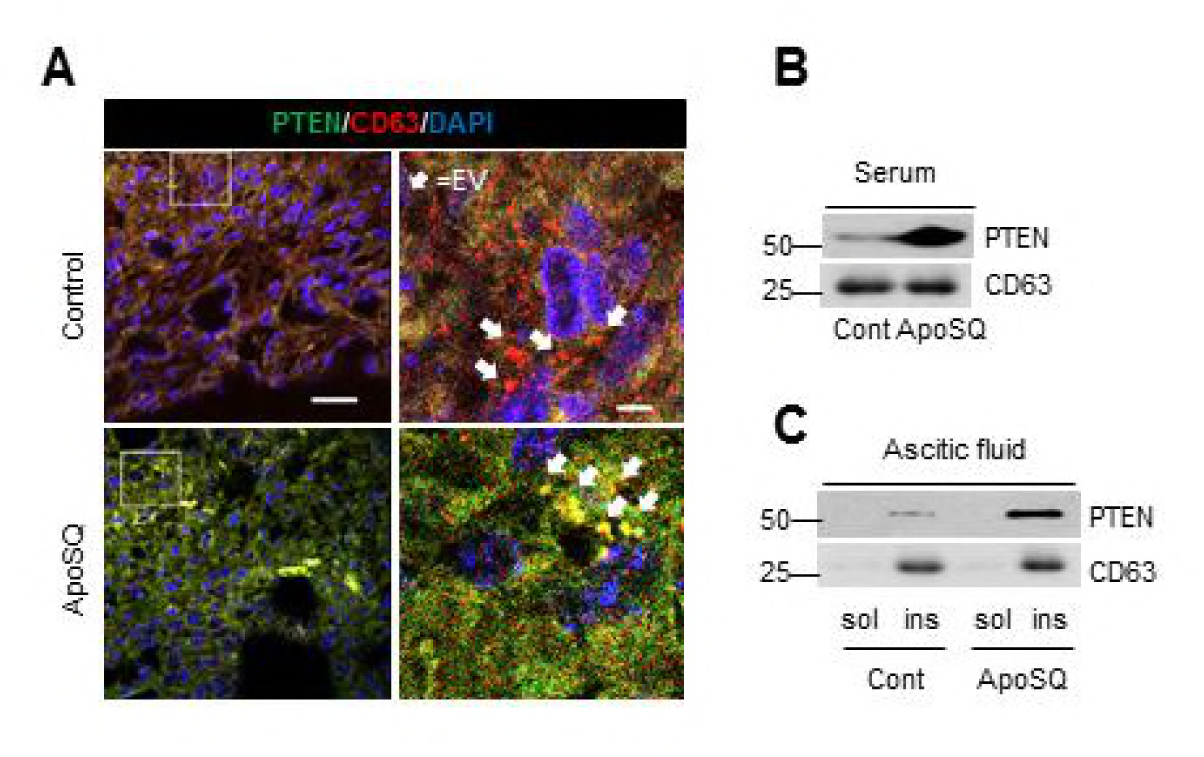
Exosomal PTEN release in tumor tissue and PTEN levels in body fluid following apoptotic cancer cell injection in mice. (A-C) Apoptotic 344SQ cells (ApoSQ) were subcutaneously (SQ) injected in the skin lesion 2 days after subcutaneous injection of 344SQ cells into syngeneic (129/Sν) mice. Mice were necropsied 6 weeks later. (A) Confocal images of primary tumor sections stained with anti-PTEN (green) or anti-CD63 (red), and the DNA-binding dye DAPI. (A *right*) Enlarged ROIs from white squares on the low magnification images of A *left*. Arrows indicate the localization of extracellular vesicles (EV) in tumor microenvironment. Scale bars: 20 μm (*left*) and 1 μm (*right*). Immunoblot analysis of PTEN and CD63 in serum using Total Exosome Isolation kit (B) and in the sol and ins fractions after ultracentrifugation of ascitic fluid (C). Data are representative images from five mice per group (A-C)

To confirm PPARγ-dependent PTEN expression and concurrently mediating ant-metastasis effect, the PPARγ antagonist GW9662 (1mg/kg/d) was i.p. administered for 4 weeks, beginning one day before ApoSQ injection. GW9662 treatment reversed reduction of metastatic incidence by ApoSQ injection (Figure 9A). Interestingly, the enhancement of *PTEN* and *CD36* and the reduction of *Snai1* and *Zeb1* mRNA expression in tumor tissue by ApoSQ injection were reversed by GW9662 treatment (Figure 9B). Moreover, ApoSQ-induced enhancement of PTEN protein expression and reduction of Akt phosphorylation were also reversed by GW9662 treatment (Figure 9C). This inhibitor administered with buffer had no effects.

**Figure 9.**
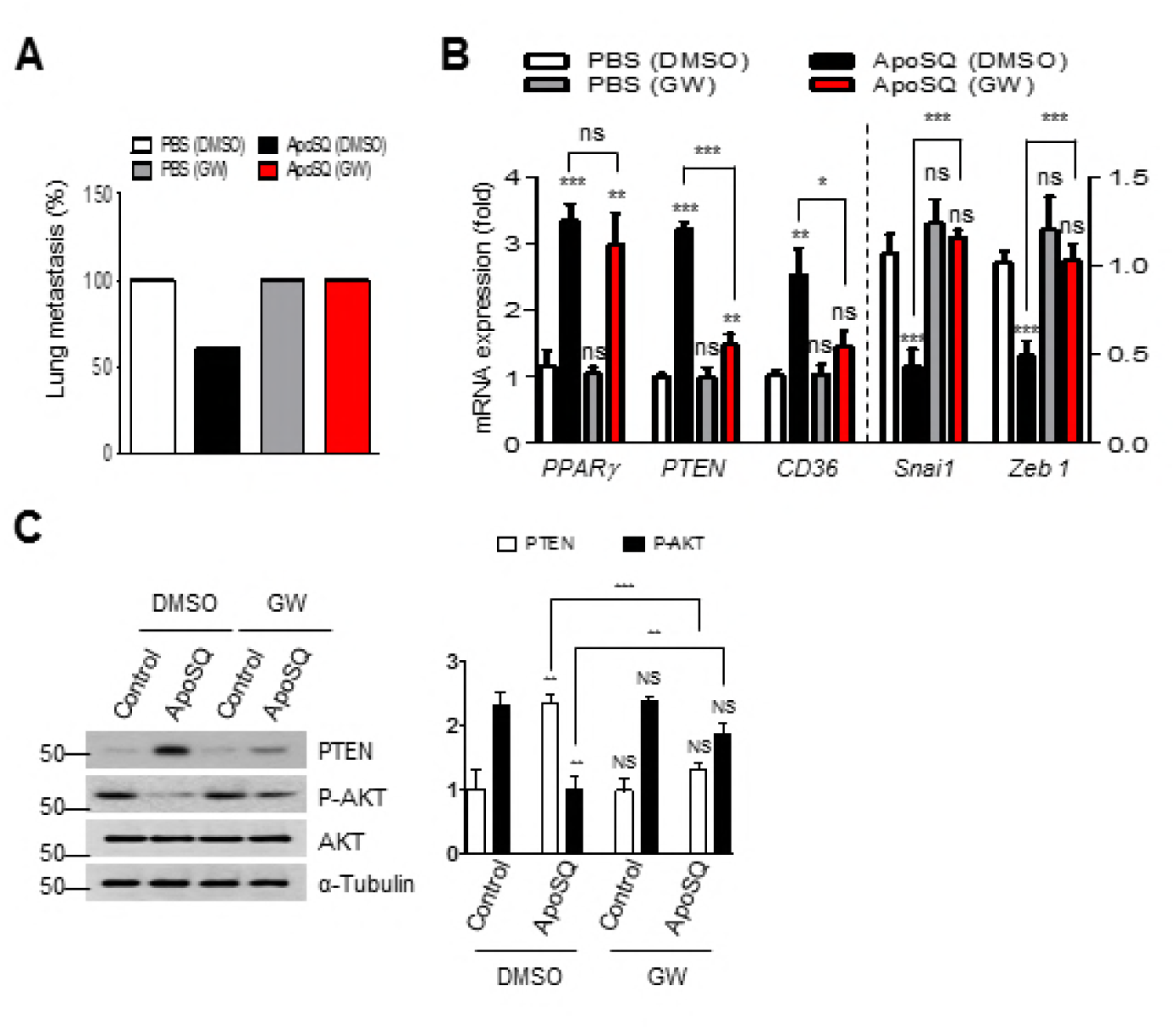
PPARγ-dependent anti-metastatic effects of apoptotic cell injection in mice. (A-C) Where indicated, GW9662 (1 mg/kg/day, i.p.) or its vehicle (Veh; 2% DMSO in saline) was administered into the left flank one day before SQ injection of ApoSQ in the right skin lesion (n=5 per group). Mice were necropsied 4 weeks after 344SQ cell injection. (A) Lung metastasis incidence (%). qPCR analysis (B) and immunoblot analysis of indicated protein expression (C) in primary tumors. \**P* < 0.05, \*\**P* < 0.01 and \*\*\**P* < 0.001 (Student’s *t* test). Data are representative images from five mice per group (C *left*) or from independent experiments with five mice per group (mean ± s.e.m. in B and C).

## Discussion

EMT activation has previously been proposed as the critical mechanism in malignant phenotype acquisition by epithelial cancer cells (Thiery, 2002); we now propose that the interaction between macrophages and apoptotic cancer cells could provide an anti-cancer microenvironment to inhibit EMT and the multistep process of cancer cell dissemination. Our in vitro data demonstrate that ApoSQ-exposed CM from RAW cells and primary mouse BMDMs inhibited TGF-β1-induced EMT in 344SQ cells. In addition to murine lung adenocarcinoma cells, the interaction of macrophages with various types of apoptotic human cancer cells, such as non-small cell lung, breast, colon, and prostate cancer cells, but not necrotic cells, results in the inhibition of EMT marker changes. These data clearly suggest that this anti-EMT effect is universal and specific. Similar to wild-type mouse and human macrophages, CM from M2-like BMDMs sharing properties TAM phenotype (Sica et al., 2006), and blood MDMs of lung adenocarcinoma patients exposed to apoptotic cancer cells, show anti-EMT effects. These data suggest that macrophages under normal or cancer circumstances may consistently have the ability to prevent EMT in response to apoptotic cancer cells.

Our data suggest that downregulation of Smad-independent signaling, including p38 MAP kinase and Akt pathways, by ApoSQ-exposed CM inactivates transcription factors that bind to the *Snai1/2, Zeb1/2, and Twist1* promoters in the tumor microenvironment. We found that apoptotic cancer cell-exposed CM inhibits migration and invasion, as well as cancer stem-like phenotype acquisition, in cancer cells. Moreover, anti-invasive effect of the ApoSQ-exposed CM was confirmed using 3D Matrigel culture. These findings provide the new insight that macrophages exposed to apoptotic cancer cells might create a tumor microenvironment preventing metastatic processes.

This is the first report of PPARγ activity-dependent PTEN induction in macrophages exposed to apoptotic, but not necrotic, cancer cells. Recently, two different groups demonstrated that PTEN secreted via exosome formation, or PTEN-Long via an unknown mechanism, can be internalized by recipient cells (Hopkins et al., 2013, Putz et al., 2012). Surprisingly, we found enhanced canonical PTEN expression in PTEN immunoprecipitates from ApoSQ-exposed CM, and confirmed the secretion of this protein via exosome formation from multivesicular bodies. PTEN secretion in exosomes by human macrophages overexpressing GFP-PTEN in response to ApoA was also verified. Moreover, PTEN-bearing exosomes appear to be uptaken by recipient cancer cells. The way individual vesicles interact with recipient cells is still not known, and has been proposed to involve binding at the cell surface via specific receptors, internalization by a variety of endocytic pathways or micropinocytosis, and/or fusion with plasma membrane or with the limiting membrane of internal compartments (Thery, 2011). Accordingly, PTEN abundance in recipient 344SQ cells was enhanced immediately, and remained until 24 h after treatment with ApoSQ-exposed CM, without changes in PTEN mRNA expression; Akt phosphorylation decreased reciprocally. Interestingly, ApoSQ-exposed CM from PTEN knockdown RAW cells failed to enhance PTEN abundance and reduce TGFβ-induced p38 MAP kinase and Akt phosphorylation. Taken together, these data indicate that enhanced PTEN protein levels in 344SQ cells do not originate from early-phase transcriptional induction, but from its internalization into recipient cells with intact lipid and possibly protein phosphatase activity (Tibarewal et al., 2012).

PTEN functions in a spatially restricted manner, which may explain its involvement in forming PIP3 gradients, necessary for generating and/or sustaining cell polarity in epithelial tissues (Martin-Belmonte et al., 2007). Accumulating evidence indicates that loss of cellular polarity and tissue architecture can drive tumor progression (Wodarz and Nathke, 2007). We observed enhanced PTEN recruitment to the cancer cell plasma membrane 12 h after the addition of ApoSQ-exposed CM, and concomitantly reduced Akt phosphorylation in the plasma membrane-enriched fraction. Moreover, our data from the experiments using PTEN knockdown in RAW cells or PTEN signaling inhibition in 344SQ cells highlight that signaling through internalized PTEN mediates prolonged anti-EMT and anti-invasion effects, controlling early cell polarity and integrity by preventing the dissolution of cell-cell contacts.

On the other hand, based on the prolonged effects of restored *PTEN* mRNA expression over 48 h following ApoSQ-exposed CM treatment, we propose that PTEN signaling in recipient 344SQ cells may originate from a different source, although the PTEN internalization rate, half-life, and stability of internalized PTEN were not estimated (Vazquez et al., 2000). Notably, with regard to PPARγ activation over 72 h after treatment with ApoSQ-exposed CM, 344SQ cells demonstrated a striking resemblance to cells with *PTEN* mRNA expression, with the dependence on PPARγ activity. These data support the novel insight that enhanced late-phase *PTEN* mRNA expression may be induced mainly through PPARγ-dependent transcriptional upregulation in recipient cancer cells.

We observed enhanced secretion of PPARγ ligands in ApoSQ-exposed CM, but not in viable or necrotic 344SQ-exposed CM. Interestingly, ApoSQ-exposed CM deficient in these PPARγ ligands partially failed to reverse the reduction of *PPARγ* mRNA and activity, and PTEN mRNA and protein levels, in recipient 344SQ cells, and consequently, EMT under TGF-β1 stimulation was not effectively prevented. These data suggest that ligand-dependent PPARγ/PTEN signaling in 344SQ cells also mediates the anti-EMT effects of ApoSQ-exposed CM. Supporting this hypothesis, the exogenous addition of these lipid mediators into 344SQ cells partially reversed TGFβ1-induced reduction of *PPARγ* mRNA expression and activation, and PTEN mRNA and protein expression, and concomitantly inhibited EMT. Unlike ApoSQ-exposed CM, these mediators did not affect early-phase TGFβ1-induced Akt and p38 MAP kinase phosphorylation, indicating that their anti-EMT effect bears no relation to early signaling events.

We identified the original sources of PTEN signaling in 344SQ cells over time after ApoSQ-exposed CM treatment: internalized PTEN (predominant in the early phase), and ligand-dependent PPARγ signaling (predominant in the late phase). Furthermore, cell polarity maintenance in the early phase after treatment with the CM is attributed primarily to internalized PTEN, as inhibiting PPARγ activity in RAW cells suppresses retaining polarity of 344SQ cells at 12 h after ApoSQ-exposed CM treatment. However, inhibited PPARγ ligand production in macrophages could not suppress this early effect of ApoSQ-exposed CM.

Increasing evidence indicates that PTEN loss triggers EMT in many cancer cell types, and consequently promotes invasion and metastasis in various cancers (Wang et al., 2015, Mulholland et al., 2012, Bowen et al., 2009). Intraperitoneally injected PTEN-Long (a translational variant of PTEN) led to tumor regression in several xenograft models, dependent on PTEN-Long phosphatase activity (Hopkins et al., 2013). Notably, PTEN deletion in stromal fibroblasts accelerated the initiation, progression, and malignant transformation of mammary epithelial tumors (Trimboli et al., 2009). In the present study, a single administration of ApoSQ around the lesion two days after 344SQ cell injection into syngeneic mice diminished number of metastatic nodules and lung metastasis incidence after 6 weeks, but caused no significant changes in primary tumor size, indicating the anti-metastatic effect of ApoSQ injection *in vivo*. However, some mice treated with ApoSQ developed lung metastases after 6 weeks of treatment with 344SQ cells. ApoSQ use should be modulated; more injections, modified injection timing, combined therapy with efferocytosis-stimulating agents, or PTEN-bearing exosome therapy with PPARγ ligands might be needed. Importantly, a single ApoSQ injection leads to enhanced induction of *PPARγ* and PTEN mRNA and protein expression, and a reciprocal reduction of phosphorylated Akt, as well as mRNA levels of *Snai1 and Zeb1*, within primary tumor tissue. Importantly, confocal microscope analysis confirms apparent PPARγ and PTEN induction in tumor cells and infiltrating macrophages. Based on our *in vitro* findings, these data suggest that *in vivo* treatment with apoptotic cancer cells creates a tumor microenvironment antagonizing metastasis via PPARγ and PTEN signaling in both macrophages and cancer cells. Along with PTEN expression, the PPARγ target gene CD36, an efferocytic surface receptor (Korns et al., 2011), was also enhanced by ApoSQ injection, suggesting that prolonged PPARγ expression in tumor-infiltrating macrophages might reinforce apoptotic cell recognition and clearance via CD36 transactivation, and prevent a defect in the ability of macrophages to clear dying cancer cells in the tumor microenvironment (Yoon et al., 2015, Mukundan et al., 2009). Not surprisingly, we found increases in PPARγ, PTEN, and CD36 mRNA or protein expression in TAM isolated from primary tumor after ApoSQ injection. In addition, enhanced production of PPARγ ligands, such as 15-HETE, lipoxin A4 and 15d-PGJ_2_, was detected in culture media of TAM. These *in vivo and ex vivo* data suggest that early injection of apoptotic cancer cells results in shifting TAM phenotype to anti-metastatic nature, leading to PPARγ-dependent PTEN and its ligand production. Enhanced PPARγ and PTEN expression in F4/80 positive macrophages in lesional skin sections within 24 h after ApoSQ injection suggests that the *in vivo* relevance of the early macrophage response to apoptotic cancer cells is to induce PPARγ-dependent PTEN production.

Similar to *in vitro* findings, detection of exosomal PTEN in the tumor sections, and body fluids, such as serum and ascitic fluid, after ApoSQ injection raised the possibility that local secretion of PTEN-bearing exosome can deliver their cargo into adjacent recipient cells, circulating in body fluids beyond tumor microenvironment. TAM may be a possible and strong candidate for exosomal PTEN sources, based on our *in vitro* and *in vivo* findings. Using PPARγ antagonist GW9662, we confirmed the new concept of PPARγ-dependent PTEN signaling as well as reduction of *Snai1* and *Zeb1* mRNA expression and concurrently mediating anti-metastasis effect.

In summary, we propose that PTEN secretion in exosomes from macrophages exposed to apoptotic cancer cells can be internalized into recipient cancer cells, inhibiting cell polarity disruption, EMT and invasion. In addition, secretion of the ligands 15-HETE, lipoxin A4, and 15d-PGJ_2_ from macrophages in response to apoptotic cancer cells may reinforce tissue homeostasis and defense against cancer metastasis via enhanced PPARγ/PTEN signaling in cancer cells. Thus, programming macrophages by apoptotic cancer cells might create a tumor microenvironment antagonizing cancer progression and metastasis via prolonged PPARγ/PTEN signaling in both TAM and tumor cells. Importantly, our studies provide new opportunities to develop apoptotic cell therapy as an effective anti-metastasis tool in a variety of experimental and clinical settings.

## Materials and Methods

### Reagents

GW9662 (#70785) and PD146176 (#10010518) were from Cayman Chemical. SC79 (SML0749) and SF1670 (SML0684) was from Sigma-Aldrich. TGF-β1 (240-B-010) and IL-4 (404-ML) were from R&D Systems. 15-HETE and 15d-PGJ_2_ were from Enzo Life Sciences. Lipoxin A4 came from Neogene. Mouse IgG Blocking Reagent (MKB-2213) was from Vector Laboratories.

### Antibodies

The antibodies used for the Western blotting and immunofluorescence are listed in Supplementary table 1.

### Cell lines, primary cells, and culture

Murine RAW 264.7 cells and human cancer cell lines were obtained from ATCC (American Type Culture Collection). 344SQ cells (gift from Dr. Kurie) (Gibbons et al., 2009) and various human cancer cell lines [A549 (lung), MDA-MB-231 (breast), COLO320HSR (colon) and PC3 (prostate)] were maintained in RPMI 1640 (HyClone™, GE Healthcare) containing 10% fetal bovine serum (FBS) and 1% penicillin/streptomycin. RAW 264.7 cells were grown in DMEM, (Gibco™, Thermo Fisher Scientific) supplemented with 10% FBS, and 1% penicillin/streptomycin. Bone marrow cells from C57BL/6 mice were cultured in DMEM supplemented with 10% FBS and 20% L929 supernatant (BMDM medium) for 7 days. For differentiation of M2-like cells, BMDMs were grown in 50 ng/ml IL-4-containing BMDM media for 3 days (McWhorter et al., 2013).

### Conditioned medium

Murine macrophages (RAW, BMDM and M2-like cells) or human blood MDM were plated at 5 × 10^5^ cells/ml and grown in suitable medium (refer to Cell cultures) at 37°C and 5% CO_2_. After overnight incubation, the cells were then serum-starved with X-VIVO 10 medium (04-380Q, Lonza) for 24 h before cell stimulations. For the stimulation, the culture medium was replaced with X-VIVO 10 containing apoptotic or necrotic cancer cells (1.5 × 10^6^ cells/ml). After 24 h, supernatants were harvested by centrifugation and used as the conditioned medium for stimulation of target cancer epithelial cells (5 × 10^5^ cells/ml).

### Blood samples from patients

Lung cancer patients and healthy controls were included in the study after informed consent under protocols approved by the Institutional Review Board of Ewha Womans University, School of Medicine. A total of three health control (one male, two females) and three non-small cell lung cancer patients without anticancer drugs (one male, two females) were used in the experiments depicted in Figure 1C, 2C and Supplementary figure 1J,K. Human monocytes were collected from 20 ml blood by Ficoll-Histopaque density gradient centrifugation (Repnik et al., 2003). Purified monocytes were grown in RPMI containing 10% human AB serum for 8 days. The confirmation of the monocyte differentiation into macrophages (MDM) was done by confocal microscopy with anti-F4/80.

### Induction of cell death

Cancer epithelial cell lines were exposed to ultraviolet irradiation at 254 nm for 10 min followed by incubation in RPMI-1640 with 10% FBS for 2 h at 37°C and 5% CO_2_. Evaluation of nuclear morphology using light microscopy on Wright-Giemsa-stained samples indicated that the irradiated cells were approximately apoptotic (Byun et al., 2014). Lysed (necrotic) cancer cells were obtained by multiple freeze-thaw cycles (Fadok et al., 2001). Apoptosis and necrosis were confirmed by Annexin V-FITC/propidium iodide (BD Biosciences, San Jose, CA) staining followed by flow cytometric analysis on a FACSCalibur system (BD Biosciences) (Byun et al., 2014).

### Immunoprecipitation

60 ml conditioned medium was prepared from BMDMs that had been non- or apoptotic 344SQ cell-stimulated at 1 × 10^6^ per ml. In the case of unconventional secretion of PTEN (Chua et al., 2014), the medium was diluted 1:1 with 2X exosome lysis buffer (4% SDS, 2% Triton-X100, 0.1 M Tris pH 7.4 and 2X protease inhibitors) (de Jong et al., 2012) and lysed for 1 hour at 4°C. The resulting medium-lysis buffer mixture was filtered through 0.22 micron filter (Macherey-Nagel) and divided into 15 ml vials. 25 μl of anti-PTEN (138G6, Cell Signaling) was added to a mixture vial and reacted for 4 h with rotation at 4°C. Immunocomplexes were then precipitated with 100 μl (50% slurry) of Protein A/G Sepharose (BioVision Inc). Pull-down beads prebound immunocomplex were added into the new tube and incubated for 4 h with rotation at 4°C repeatedly for the rest of mixture vials, and washed for times with IP wash buffer (25 mM HEPES pH7.4, 1 M Nacl, 1 mM EDTA, 0.5 % Triton X-100). To avoid overlapping with IgG Heavy chains, the Western blotting for PTEN immunoprecipitates was performed in non-reducing conditions.

### Western blotting

Standard western blottings were performed using whole cell extracts, (in)soluble fractionates from conditioned media, or immunoprecipitates. The information of antibodies was included in Supplementary table 1.

### Real-time quantitative PCR (qPCR)

mRNAs were extracted from cells grown to 80% confluence in triplicate on 6-well plates in the experimental conditions and quantified using Real-Time PCR System (Applied Biosystems, Step One Plus). See Supplementary table 2 for primer sequences of target genes.

### Migration and invasion assays

Cell migration and invasion were tested using Transwell chambers (Corning Inc) coated with 10 μg/ml fibronectin and 200 μg/ml Matrigel matrix according to the manufacturer`s instruction, respectively. In brief, pre-incubated cancer cells (5 × 10^4^ cells/well for the migration assay and 2 × 10^5^ cells/well for the invasion assay) in the conditioned medium from macrophages in the absence or presence of 10 ng/ml TGF-β1 were plated in replicate wells in serum-free RPMI in the upper chambers and in RPMI 1640 supplemented with 10% FBS placed in the bottom wells at 37°C for 16 h migration time or 24 h invasion time. After fixation in 4% paraformaldehyde, the nonmigrated or noninvaded cells on the upper surface of the membrane were scraped off with a cotton swap. The cells on the lower surface were stained using 0.1% crystal violet, and washed with distilled water. Three random microscopic fields (10X magnification) were photographed and counted.

### 3D cell culture on matrigel

Standard 3D culture was performed as described previously (Debnath et al., 2003). Briefly, a single-cell suspension containing 5000 cells/well was plated on top layer of the solidified Growth Factor Reduced Matrigel (Corning Inc) in 8 well plate. The cells in RPMI 1640 with 10% FBS and 2% Matrigel were incubated and the medium was changed every two or three days for a week. After incubation, the cells were treated with the indicated conditioned medium containing TGF-β1 (10 ng/ml) and 2% Matrigel, and then grown for 3 days. Phase contrast images were taken using Eclipse TE-300 microscope (Nikon).

### Immunofluorescence

344SQ cells grown on glass coverslips until confluent were fixed with 4% paraformaldehyde (PFA) solution for 8 min at room temperature (RT). For staining of 344SQ acini in Matrigel 3-D culture, paraffin-embedded tumor or frozen skin tissues, formalin fixation was performed at RT for 30 minutes and IF-Wash buffer (0.05% NaN3, 0.1% BSA, 0.2% Triton X-100 and 0.05% Tween-20 in PBS) was used. After fixation, samples were washed three times with wash buffers for 5 min each and permeabilized with 0.5% Triton X-100 in PBS at RT for 5 min. 5% bovine serum albumin in PBS and –containing Mouse IgG Blocking Reagent were used for ICC and IHC, respectively. Subsequently, all slides stained with antibodies were mounted with Vectashield Mounting Medium containing DAPI (Vector Laboratories, Inc), and imaged with a confocal microscope (LSM 800, Carl Zeiss). The antibody information for sources or dilution ratios is described in Supplementary table 1.

### Membrane fractionation

Plasma membrane fractions were prepared from each 150 mm dish of 344SQ cells that had been treated with the indicated conditions as described in Figure 3A using Mem-PER™ Eukaryotic Membrane Protein Extraction Reagent Kit (89826, Thermo Fisher Scientific) following the manufacturer’s protocols.

### siRNA transfection

RAW 264.7 cells were transiently transfected with siRNA specifically targeting or control siRNA (SN-1003_AccuTarget™ Negative Control; Bioneer Inc) at both 100 nM final concentrations using GeneSilencer^Ⓡ^siRNA Transfection Reagent (Genlantis Inc) according to the manufacturer’s instruction. After overnight transfection, the cells were cultured in suitable medium for 24 h and stimulated with apoptotic 344SQ cells. The siRNA sequences used for targeting genes were as follows (gene: sense, antisense). PPARγ: 5′-AGUAUGGUGUCCAUGAGAU-3′, 5′-AUCUCAUGGACACCAUACU-3′; PTEN: 5′-CAGGAAUGAACCAUCUACA-3′, 5′-UGUAGAUGGUUCAUUCCUG-3′; L-PGDS: 5′-CAACUAUGACGAGUACGCUCUGCUA-3′, 5′-GACUUCCGCAUGGCCACCCUCUACA-3′.

### Transmission electron microscopy

Conventional TEM sample preparation was done with RAW264.7 macrophages stimulated by apoptotic 344SQ cells. Ultra-thin sections were then imaged with a transmission electron microscope (H-7650; Hitachi) run at an accelerating voltage of 80 kV. iTEM software (Olympus) was used for image acquisition.

### Detection of PPARγ ligands

PPARγ ligands in the conditioned medium were measured using ELISA kits [15-HETE (ADI-900-051, Enzo Life Sciences), lipoxin A4 (#407010, Neogene), PGD_2_ (MBS703802, MyBioSource) and 15d-PGJ_2_ (ADI-900-023, Enzo Life Sciences)] according to the manufacturer’s protocols.

### PPARγ activity assay

PPARγ activity was determined in nuclear extracts (8 g) from pharmacological inhibitor-pretreated or siRNA-transfected RAW264.7 or 344SQ cells using a TransAM™ PPARγ Transcription Factor Assay kit (40196, Activ Motif Inc) according to the manufacturer’s instructions.

### Exosome purification

Exosomes were isolated from cell culture media by differential centrifugation as described previously (Putz et al., 2015). In brief, supernatant from RAW264.7 cells exposed to apoptotic or necrotic 344SQ cells was subjected to serial centrifugation of 200 g, 20,000 g and 100,000 g for clearance of dead cells, cell debris and non-exosomal fraction. After washing the exosome pellet with ice-cold PBS containing protease inhibitors, the ultracentrifugation (Optima L-100K, Beckman Coulter Inc., Brea, CA, USA) was repeated for 70 min at 100,000 g to get rid of contaminating proteins. The exosome pellet was then resuspended in RIPA buffer containing protease inhibitors. For isolation of exosomes from mouse serum, exosomes were precipitated using Total Exosome Isolation kit (4478360, Thermo Fisher Scientific) was used. Western blotting analysis was performed for identification of exosomal PTEN in conditioned medium or mouse body fluids with anti-CD63 or anti-PTEN.

### Generation of stable macrophages overexpressing GFP-PTEN

Standard lentiviral transduction was performed as described previously (Kim et al., 2015). In brief, HEK293T cells were co-transfected with pLV-EGFP-PTEN, packaging (psPAX2) and envelope (pCMV-VSV-G) vectors using the TransIT^Ⓡ^-LT1 Transfection Reagent (MIR 2300, Mirus Bio, Madison, WI, USA) according to the manufacture’s instruction, and incubated for overnight. For lentiviral transduction, the reagents were replaced with fresh media and the viral supernatants were collected every 24 h after transfection. Human macrophages (hMϕ) differentiated from THP-1 monocytes (Zhang et al., 2013) were exposed to the supernatant with 8 µg/ml polybrene for 4hr every virus collection time and the infected cells were selected using with 1 μg/ml of puromycin-containing media.

### Detection of exosomal GFP-PTEN in recipient cells

GFP-PTEN overexpressing THP-1 macrophages were seeded in ten 150 mm tissue culture dishes and grown to 70% confluency. After stimulation with apoptotic A549 cells for 24 h, conditioned media were collected and subjected to serial centrifugation (refer to exosome purification). The exosome pellet was resuspended in 200 μl serum free RPMI. Recipient cells (A549 or 344SQ, 1 × 10^5^ cells for confocal microscopy or 3 × 10^5^ cells for Western blotting analysis) were exposed to 50 μl of purified exosomes for 24 h. GFP-PTEN in recipient cell were visualized by direct fluorescence, and also detected with anti-GFP using whole cell lysates.

### Mouse experiments

The Animal Care Committee of the Ewha Medical Research Institute approved the experimental protocol. Mice were cared for and handled in accordance with the National Institute of Health (NIH) Guide for the Care and Use of Laboratory Animals. Lung cancer metastasis studies with the syngeneic tumor experiments were performed as previously described (Yang et al., 2014). In brief, syngeneic (129Sv) mice (n = 23 per group) of at least 8 weeks old were used for the syngeneic tumor experiments. 1 × 10^6^ 344SQ cells in single-cell suspension were subcutaneously injected into the right posterior flank. 2 days after the first injection, second injection was performed in a volume of 100 μl of PBS with or without 1 × 10^7^ apoptotic 344SQ cells in the same lesion. Mice were monitored daily for tumor growth and sacrificed at 6 weeks after injection. Necropsies were performed to investigate the weights of subcutaneous tumor mass, the lung metastatic status (# of nodules or incidence) and the histological evaluation of formalin fixed, paraffin-embedded, immunofluorescence-stained primary tumor. For the inhibition experiments, the selective PPARγ antagonist GW9662 (1mg/kg/d) was i.p. administered for 4 weeks, beginning one day before ApoSQ injection. Mice were necropsied 4 weeks after 344SQ cell injection. For the investigation of early activation of dermal macrophages in response to apoptotic cancer cells, 20-week-old wild type of B6129SF2/J mice were injected with 344SQ cells in the same way and killed 6 or 24 h after the second injection. Right flank skin from mouse was isolated and frozen in OCT compound for immunofluorescent staining of the lesional skin tissue.

### TAM isolation

Purification of tumor-associated macrophages (TAM) was performed as described previously (Laoui et al.). To obtain single cell suspension from lung tumors of metastatic mouse models with or without injection of ApoSQ (n = 8 per group, refer to Mouse experiments), solid fresh cancer tissues were disaggregated with tumor digestion medium containing Collagenase I, IV and DNase I. After filtration with 70 μm sterile nylon gauze, red blood cells were lysed with erythrocyte lysis buffer. Density gradient centrifugation was performed to isolate mononuclear cells from sharp interphase, and sedimented cells (mostly cancer cells) were used for Western blot analysis. From mononuclear cells washed by MACS buffer, TAM were isolated using anti-CD11b+-conjugated magnetic beads and MACS columns (Miltenyi Biotec). Cross-checking for the identification of TAM was done by confocal microscopy with anti-F4/80.

### Statistics

Comparisons between 2 mean values ± SEM (control versus experimental) were performed using the two-tailed Student’s *t* test. *P* values that are less than 0.05 are considered statistically significant. All data were analyzed using Graph Prism 5 software. (GraphPad Software Inc).

## Acknowledgements

The authors thank Dr. J.M. Kurie (University of Texas MD Anderson Cancer Center) for critical reading of the manuscript and providing 344SQ cells and syngeneic (129/Sν) mice; Dr. M.C. Baek (Kyungpook National University) for technical advice on exosome isolation. This work was supported by Basic Science Research Program through the National Research Foundation of Korea (NRF) funded by the Ministry of Science, ICT & Future Planning (2015R1A2A1A15053112 and 2010-0027945).

## Author contributions

Y.B.K. designed and performed most of the *in vitro* and *in vivo* experiments and analyzed data; Y.H.A. performed some *in vivo* experiments and offered technical advice; J.H.L. provided blood from healthy humans and lung cancer patients; J.L.K. directed and designed the study, analyzed data, and wrote the paper.

---

**Supplementary figure 1.**
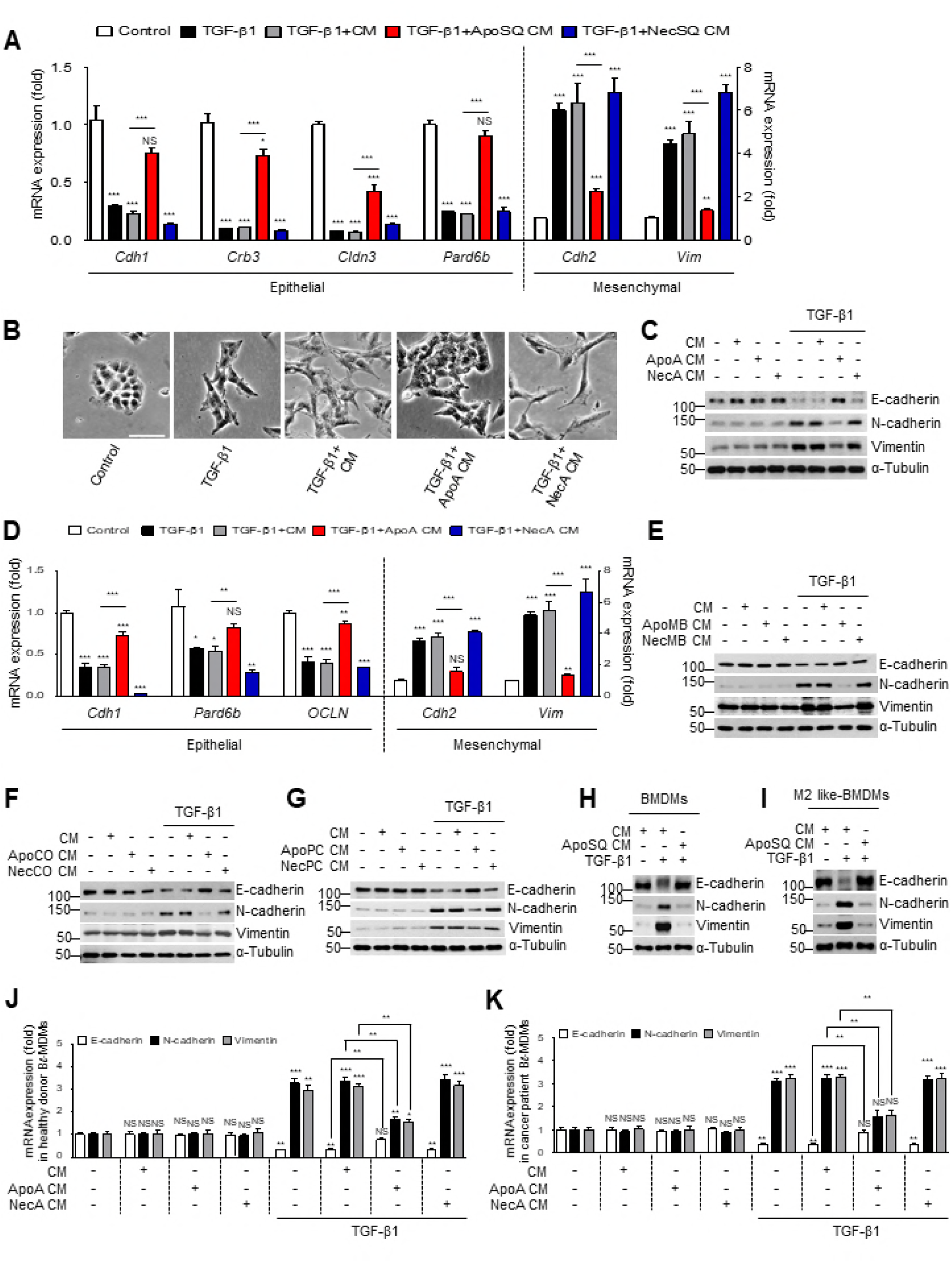
Interaction of macrophages and apoptotic cancer cells inhibits EMT in cancer cells. RAW cells were stimulated with apoptotic cancer cells, such as 344SQ cells (ApoSQ) (A), A549 (ApoA) (B-D), MDA-MB-231 (ApoMB) (E), COLO320HSR (ApoCO) (F), and PC3 (ApoPC) (G), or necrotic cancer cells (NecSQ, Nec A, NecMB, NecCO, and NecPC, in A, B-D, E, F, and G, respectively). Mouse bone marrow-derived macrophages (BMDMs) (H) or M2-like BMDMs (I) were stimulated with ApoSQ or NecSQ. Blood monocyte-derived macrophages (MDMs) from healthy donors (J) or lung cancer patients (K) were stimulated with ApoA or NecA. (A-K) After 24 h, conditioned medium (CM) was added to the corresponding cancer cells with or without TGF-β1 (10 ng/ml) for 48 h. (A and D) Real-time PCR analysis of the indicated mesenchymal markers and epithelial markers in cancer cell samples. (B) Morphological changes in the cells were examined by phase-contrast microscopy. Scale bar, 100 μm. (C and E-I) Immunoblot analysis of indicated EMT markers, and α-tubulin in total cell lysates. (J and K) Relative densinometric intensity of indicated proteins. NS, not significant; \**P* < 0.05, \*\**P* < 0.01 and \*\*\**P* < 0.001. Data are from one experiment representative of three independent experiments with similar results (B, C and E-I), or from three independent experiments (mean ± s.e.m. in A, D, J and K).

**Supplementary figure 2.**
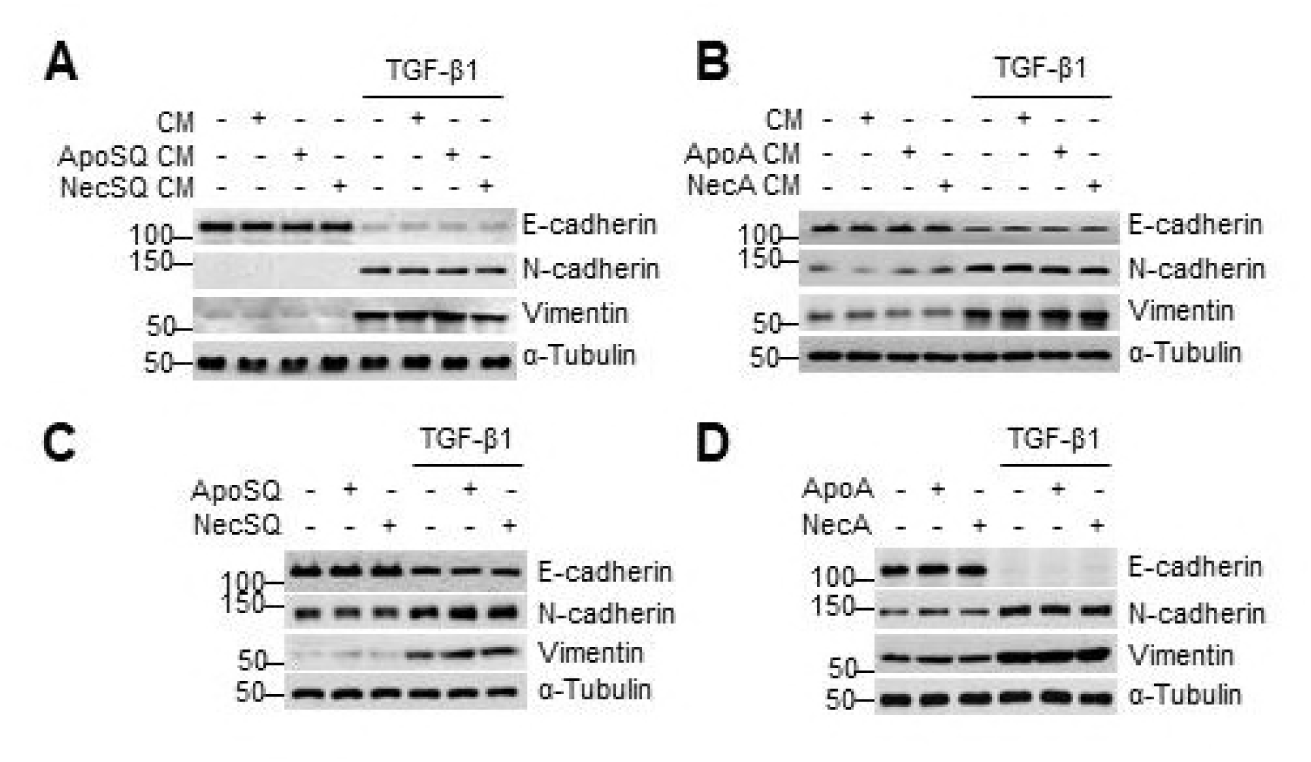
Effect of direct exposure of cancer cells to corresponding apoptotic cancer cells on TGF-β1-induced EMT. (A-D) Immunoblot analysis of EMT markers (E-cadherin, N-cadherin, and vimentin), and α-tubulin in total cell lysates. (A and B) 344SQ (SQ) (A) or A549 cells (B) were stimulated directly with ApoSQ, NecSQ, ApoA, or NecA for 24 h. Conditioned medium (CM) was added to the corresponding cancer cells in the absence or presence of TGF-β1 (10 ng/ml) for 48 h. (C and D) SQ (C) or A (D) were stimulated directly with ApoSQ, NecSQ, ApoA, or NecA in the presence of 10 ng/ml TGF-β1 for 48 h. Data are from one experiment representative of three independent experiments with similar results

**Supplementary figure 3.**
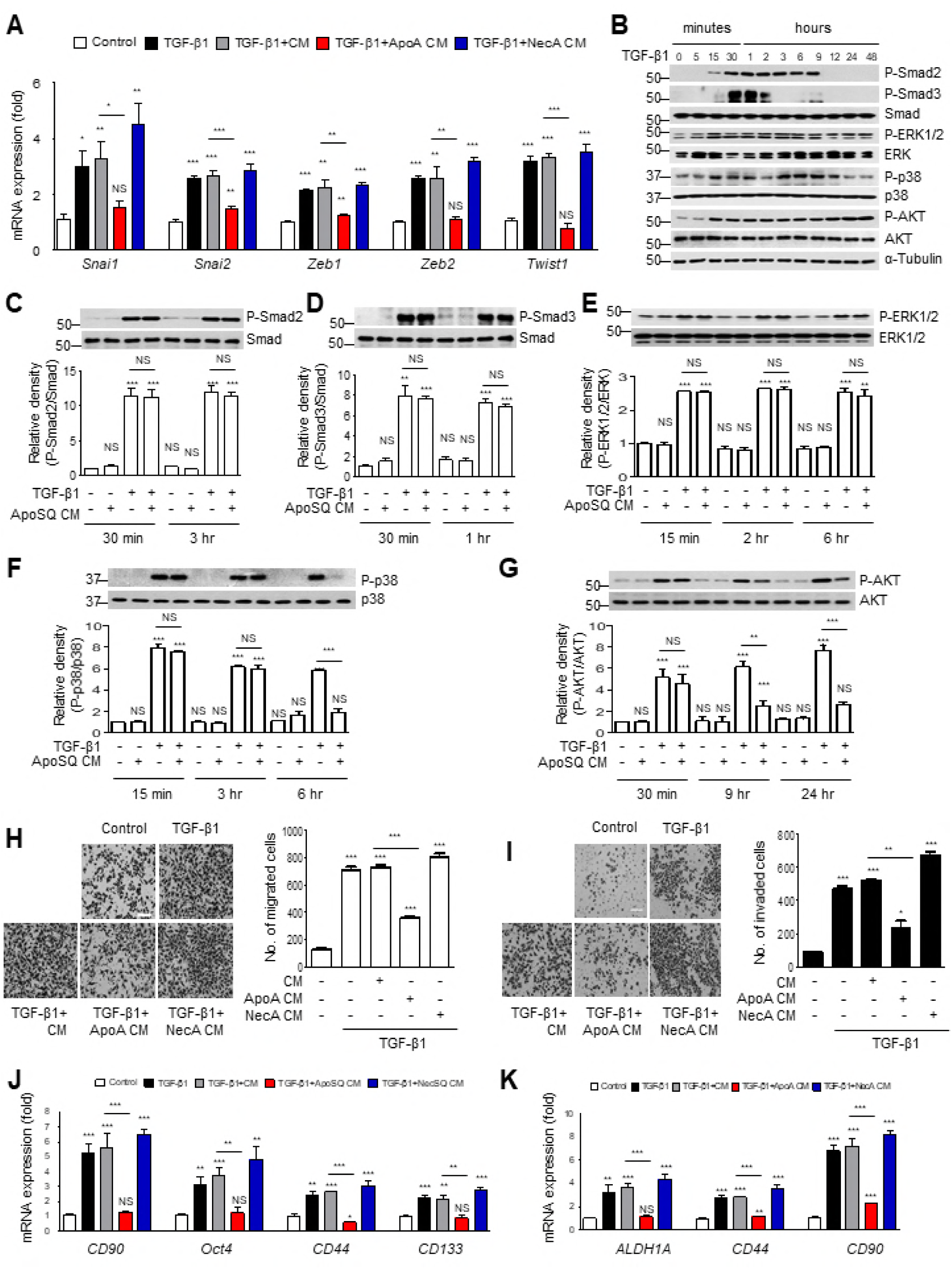
Interaction between macrophages and apoptotic cancer cells inhibits TGF-β1 signaling with progression of cancer cells. (A) A549 cells were treated with conditioned medium (CM) from RAW cells alone, or from RAW/apoptotic A549 cell (ApoA CM) or RAW/necrotic A549 cell (NecA CM) co-cultures in the presence of TGF-β1 (10 ng/ml) for 48 h. Real-time PCR analysis of *Snai1*, *Snai2*, *Zeb1*, *Zeb2,* and *Twist1* mRNAs in A549 cell samples. (B) 344SQ cells were stimulated with TGF-β1 for the indicated times. (C–G) 344SQ cells were treated with CM from RAW/apoptotic 344SQ cell (ApoSQ CM) co-cultures in the presence of TGF-β1 for the indicated times. (B–G) Immunoblot analysis of phosphorylated Smad2, phosphorylated Smad3, total Smad, phosphorylated ERK1/2, total ERK, phosphorylated p38 MAP kinase, total p38 MAP kinase, Akt phosphorylated at S473, total Akt, and α-tubulin in 344SQ cell lysates, and the relative densitometric intensity of the indicated proteins. (H, I and K) A549 cells were treated with conditioned medium (CM) from RAW cells alone, or from RAW/apoptotic A549 (ApoA CM) or RAW/necrotic A549 (NecA CM) cell co-cultures, in the presence of TGF-β1 (10 ng/ml) for 48 h. (J) 344SQ cells were treated with CM from RAW cells alone, RAW/apoptotic 344SQ cells (ApoSQ CM), or RAW/necrotic 344SQ cells (NecSQ CM) in the presence of TGF-β1 for 48 h. (H and I) The cells were visualized by phase-contrast microscopy for the analysis of migratory (H *left*) and invasive (I *left*) abilities using Fn-coated Transwell and Matrigel-coated Transwell plates, respectively. Scale bars: 100 μm. Quantification of cells that migrated (H *right*) or invaded (I *right*). Real-time PCR analysis of cancer stem-like cell markers (*CD90*, *Oct4*, *CD44*, and *CD133*) in 344SQ cells (J) and (*ALDH1A*, *CD44*, and *CD90*) in A549 cells (K) under the indicated conditions. NS: not significant; \**P* < 0.05, \*\**P* < 0.01 and \*\*\**P* < 0.001. Data are from three independent experiments (mean ± s.e.m. in A, C–G *lower panel,* J and K), from one experiment representative of three independent experiments with similar results (B, C–G *upper panel*, H and I *left*), from three fields from replicate wells (mean ± s.e.m. in H and I *right*).

**Supplementary figure 4.**
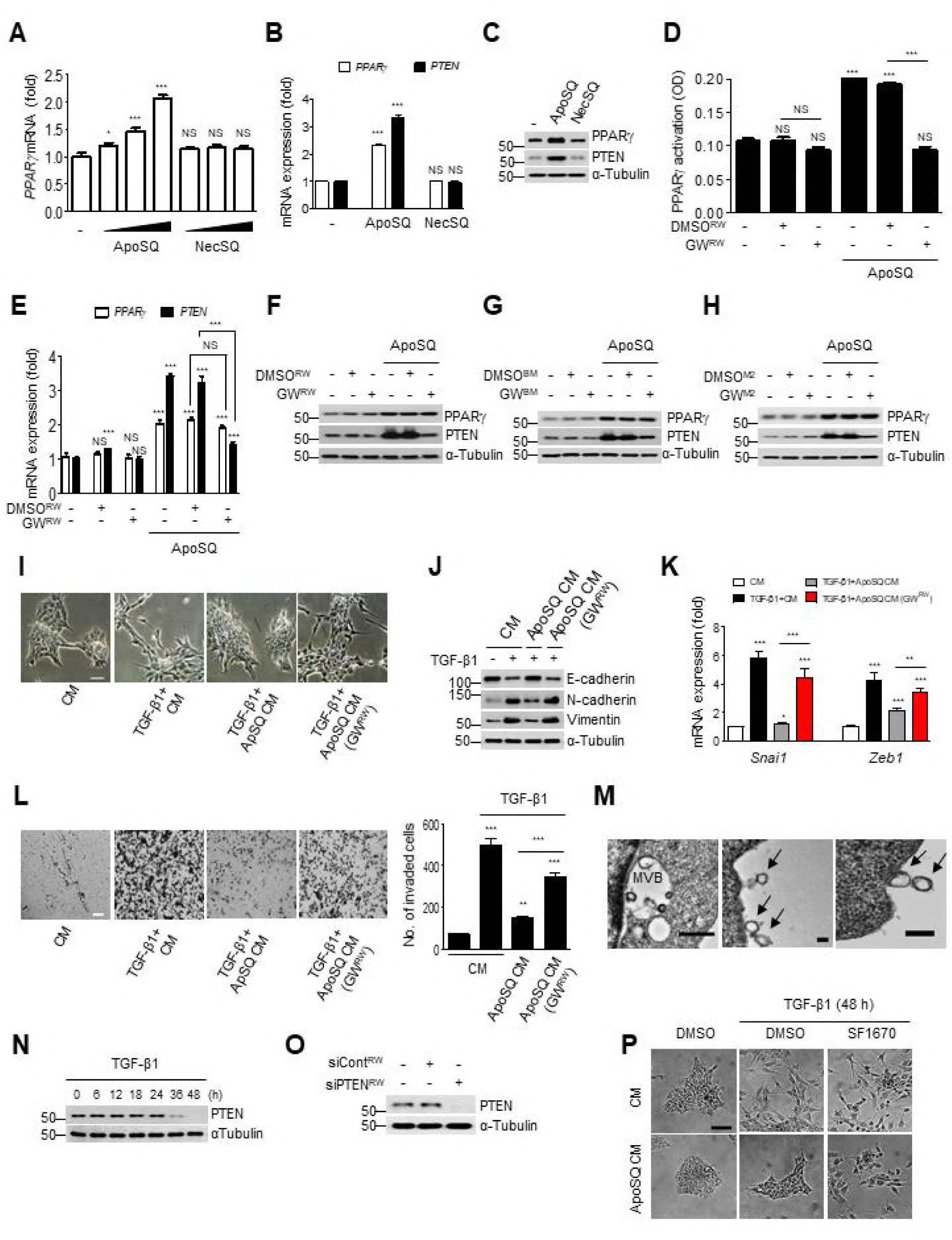
PPARγ-dependent PTEN expression in macrophages exposed to apoptotic 344SQ cells and anti-EMT effect of PPARγ/PTEN signal from macrophages in recipient cells. (A) Real-time PCR analysis of *PPARγ* mRNAs in RAW cells stimulated with apoptotic (ApoSQ; 1, 2, or 3 × 10^5^/ml) or necrotic (NecSQ; 1, 2, or 3 × 10^5^/ml) 344SQ cells for 2 h. (B) Real-time PCR analysis of *PPARγ* and *PTEN* mRNAs in RAW cells stimulated with ApoSQ cells (3 × 10^5^/ml) or NecSQ cells (3 × 10^5^/ml) for 2 h for PPARγ mRNA analysis or for 24 h for PTEN mRNA analysis. (C) Immunoblot analysis of indicated proteins in RAW cells stimulated with ApoSQ (3 × 10^5^/ml) or NecSQ (3 × 10^5^/ml) for 24 h. (D) PPARγ activity in nuclear extracts from RAW cells pretreated with GW9662 (10 μM) for 1 h before stimulation with ApoSQ (3 × 10^5^/ml) for 24 h. Real-time PCR analysis of PPARγ and PTEN mRNAs (E) and protein (F) in RAW cells pretreated with GW9662 (10 μM) for 1 h before stimulation with ApoSQ. Immunoblot analysis of indicated proteins in BMDMs (G) or M2-like BMDMs (H) pretreated with GW9662 (10 μM) for 1 h before stimulation with ApoSQ (3 × 10^5^/ml) for 24 h. (I-L) RAW cells were pretreated with GW9662 (10 μM) for 1 h before stimulation with ApoSQ cells (3 × 10^5^/ml) for 24 h. Conditioned medium (CM) was added to 344SQ cells in the presence of TGF-β1 (10 ng/ml) for 48 h. (I) Morphological changes in the cells were examined by phase-contrast microscopy. (J) Immunoblot analysis of indicated proteins in 344SQ cell lysates. (K) Real-time PCR analysis of *Snai1* and *Zeb1* mRNAs in 344SQ cells. (L) The cells were visualized by phase-contrast microscopy for the analysis of invasive ability using Matrigel-coated Transwell (*left*), and the invaded cells were quantified (*right*). Scale bars: 100 μm (I and L). (M) TEM analysis of microvesicles from apoptotic 344SQ-stimulated RAW cells. Pictures show a multivesicular body (MVB, *left panel*), and exosomes 70–100 nm in diameter released from the plasma membrane (*middle panel and right panel*). Arrows indicate exosomes and scale bars, 1 μm (left panel) and 100 nm (middle and right panel), respectively. (N) Immunoblot analysis of PTEN and α-tubulin in 344SQ cells in the presence of TGF-β1 (10 ng/ml) for the indicated times. (O) Immunoblot analysis of PTEN in RAW cells transfected with PTEN siRNA. (P) Morphological changes in the cells were examined by phase-contrast microscopy. Scale bar, 100 μm. 344SQ cells were pretreated with SF1670 (10 μM) for 1 h before treatment with CM from RAW264.7 cells or RAW264.7/apoptotic 344SQ cells (ApoSQ CM) in the presence of TGF-β1 (10 ng/ml) for 48 h. NS: not significant; \**P* < 0.05, \*\**P* < 0.01 and \*\*\**P* < 0.001. Data are from three independent experiments (mean ± s.e.m. in A-C, E and K), from one experiment representative of three independent experiments (D, F-J, L *left* -P), or from three fields from replicate wells (mean ± s.e.m in L *right*).

**Supplementary figure 5.**
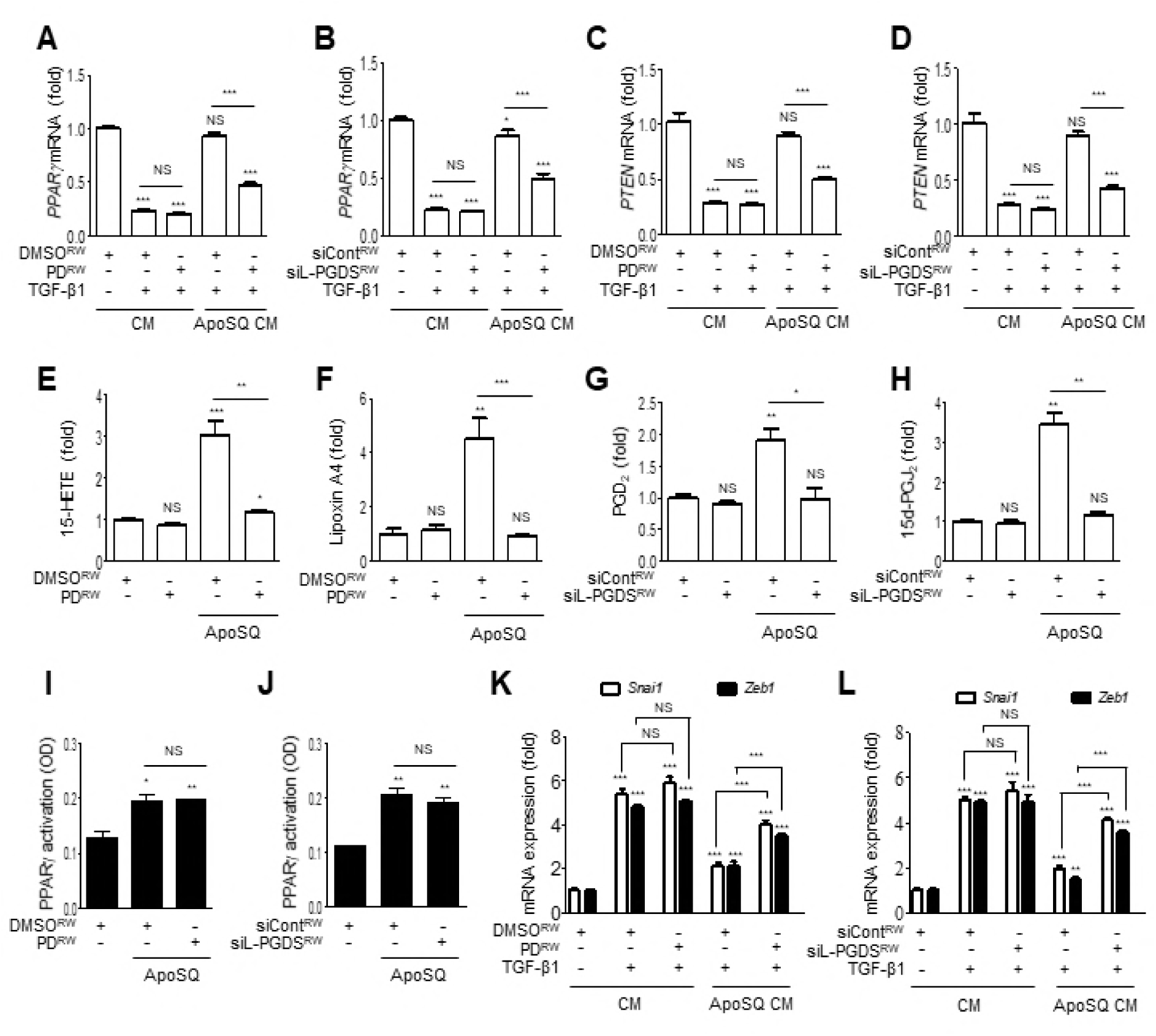
Interaction of macrophages and apoptotic cancer cells inhibits EMT regulating transcription factor via ligand-dependent PPARγ/PTEN signaling. RAW cells pretreated with PD146176 (A, C, E, F, I and K) for 1 h or transfected with siL-PGDS (B, D, G, H, J and L) for 24 h before stimulation with ApoSQ cells for 24 h. (A–D, K and L) Conditioned medium (CM) was added to 344SQ cells in the presence of TGF-β1 (10 ng/ml) for 48 h. Real-time PCR analysis of *PPARγ* (A and B), *PTEN* (C and D), *Snai1* and *Zeb1* (K and L) mRNAs in 344SQ cell samples. ELISA of 15-HETE (E), lipoxin A4 (F), PGD_2_ (G), and 15d-PGJ_2_ (H) in the CM. (I and J) PPARγ activation in nuclear extracts from RAW cells. NS: not significant; \**P* < 0.05, \*\**P* < 0.01 and \*\*\**P* < 0.001. Data are from three independent experiments (mean ± s.e.m in A–L).

**Supplementary figure 6.**
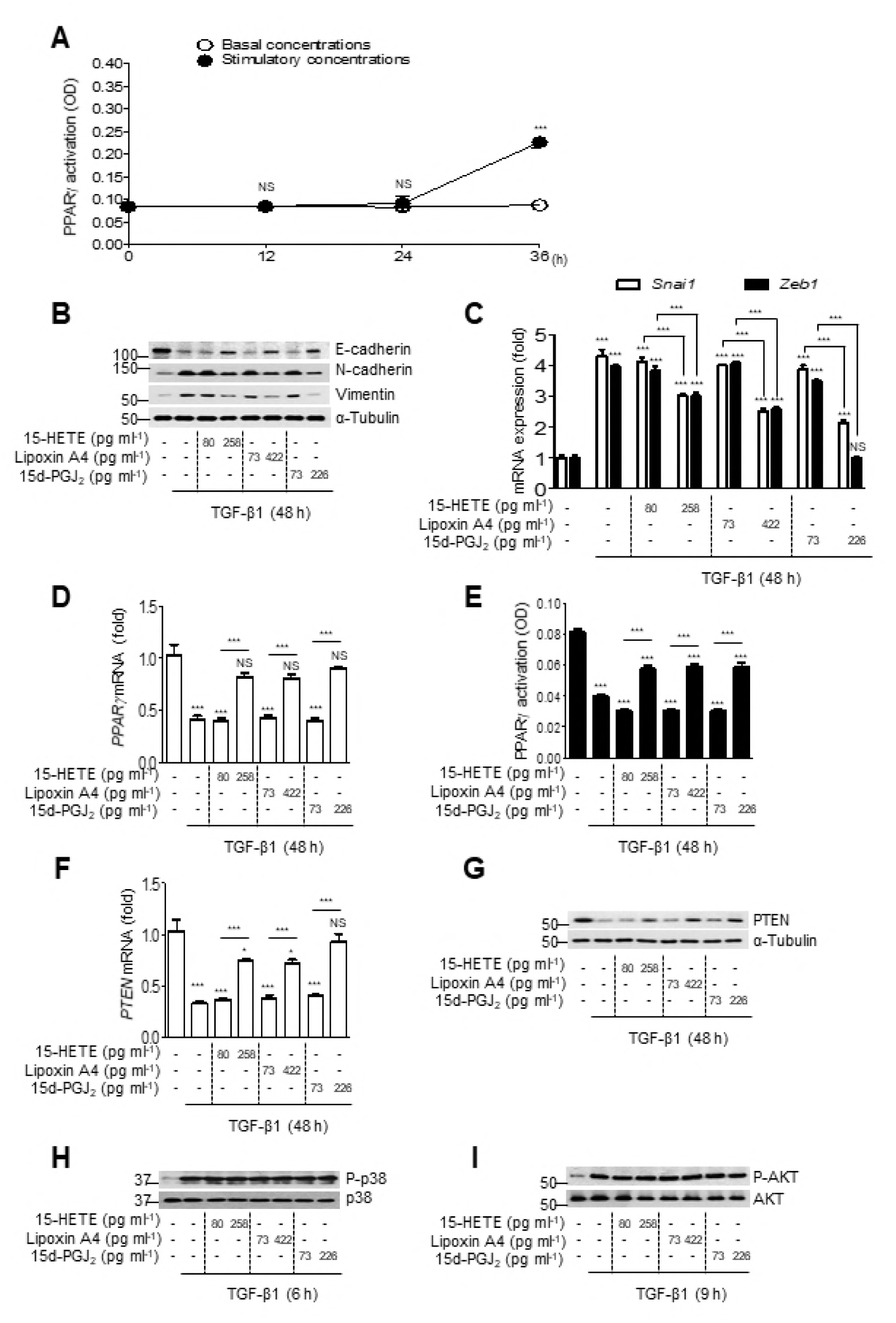
Effects of exogenous 15-HETE, lipoxin A4, and 15d-PGJ_2_ on EMT, PPARγ/PTEN signaling, and signaling. (A) Time course of PPARγ activation in 344SQ cells treated with 15-HETE (80 and 258 pg/ml), lipoxin A4 (73 and 422 pg/ml), and 15d-PGJ_2_ (73 and 226 pg/ml) all together for the indicated times. (B–I) 344SQ cells were treated individually with 15-HETE, lipoxin A4, or 15d-PGJ_2_ in the presence of TGF-β1 (10 ng/ml) for the indicated times. Immunoblot analysis of E-cadherin, N-cadherin, vimentin (B), PTEN (G), phosphorylated-p38 MAP kinase, total p38 MAP kinase (H), phosphorylated Akt, total Akt (I) and α-tubulin in total cell lysates. Real-time PCR analysis of *Snai1*, *Zeb1* (C), *PPARγ* (D), and *PTEN* (F) mRNAs in 344SQ cell samples. E, PPARγ activation in nuclear extracts from 344SQ cells. NS: not significant; \**P* < 0.05 and \*\*\**P* < 0.001. Data are from three independent experiments (mean ± s.e.m. in A and C–F), or from one experiment representative of three independent experiments (B and G–I).

**Supplementary figure 7.**
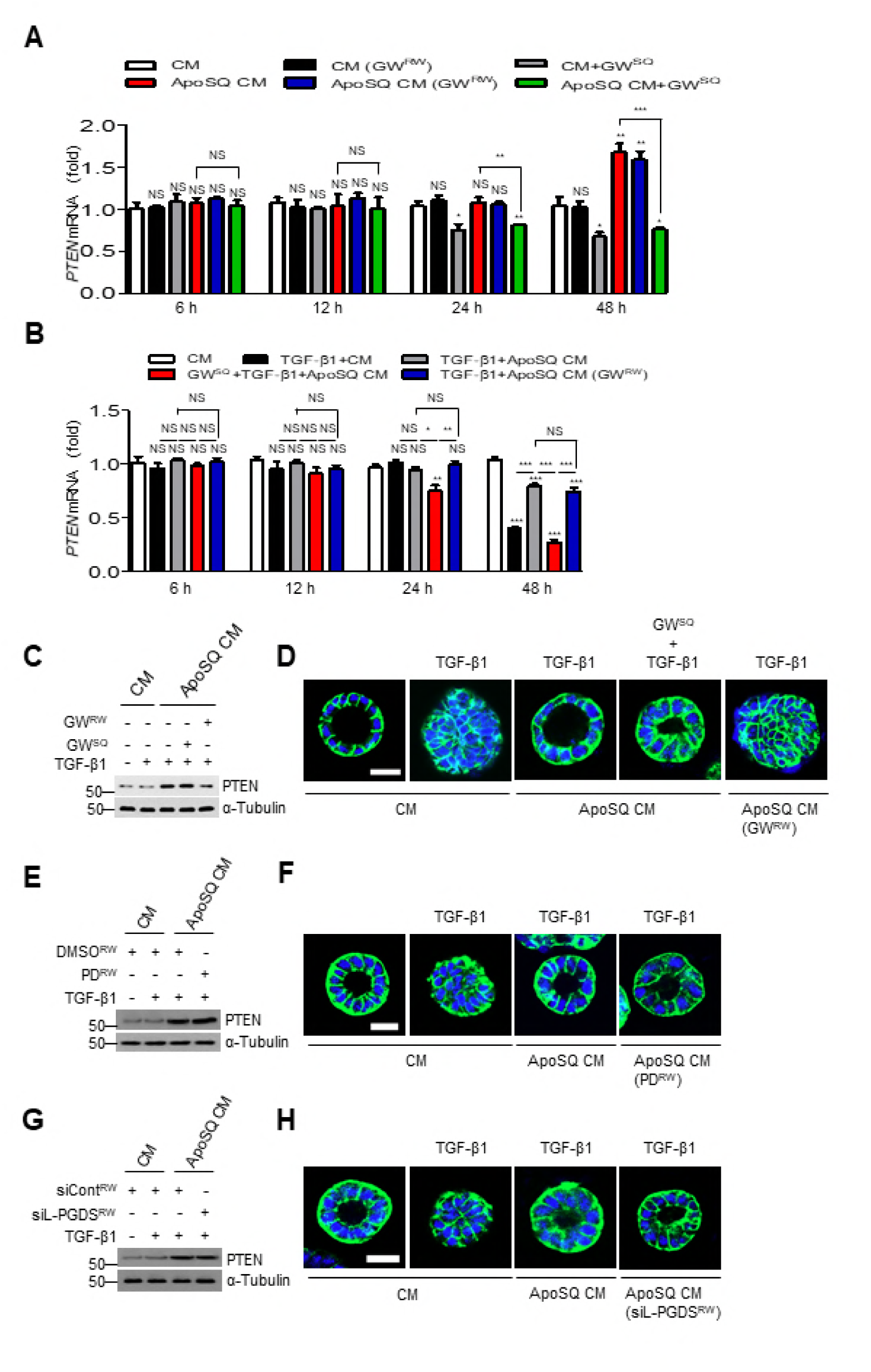
Sources of PTEN signaling in 344SQ cells are acquired from internalized PTEN and ligand-dependent PPARγ activity. (A-D) 10 μM GW9662 (GW) was used to pretreat RAW cells before treatment with apoptotic 344SQ cells (ApoSQ), or 344SQ cells before treatment with conditioned medium (CM). RAW cells were pretreated with 10 μM PD146174 (PD) for 1 h (E and F), or transfected with siRNA of lipocalin-type prostaglandin synthase (siL-PGDS) for 48 h before treatment with ApoSQ cells (G and H). CM or ApoSQ CM was added to 344SQ cells in with (A) or without TGF-β1 (10 ng/ml) for the indicated times (B) or 12 h (C, E and G). (A and B) Real-time PCR analysis of *PTEN* mRNA in 344SQ cells. (C, E and G) Immunoblot analysis of PTEN in 344SQ cell lysates. (D, F and H) Normal acini of 344SQ cells were grown in 3D Matrigel cultures containing CM with TGF-β1 for 12 h and stained with anti-β-catenin (green) and the DNA-binding dye DAPI. Scale bars: 20 μm. NS: not significant; \**P* < 0.05, \*\**P* < 0.01 and \*\*\**P* < 0.001. Data are from three independent experiments (mean ± s.e.m. in A and B) or from one experiment representative of three independent experiments with similar results (C–H).

**Supplementary figure 8.**
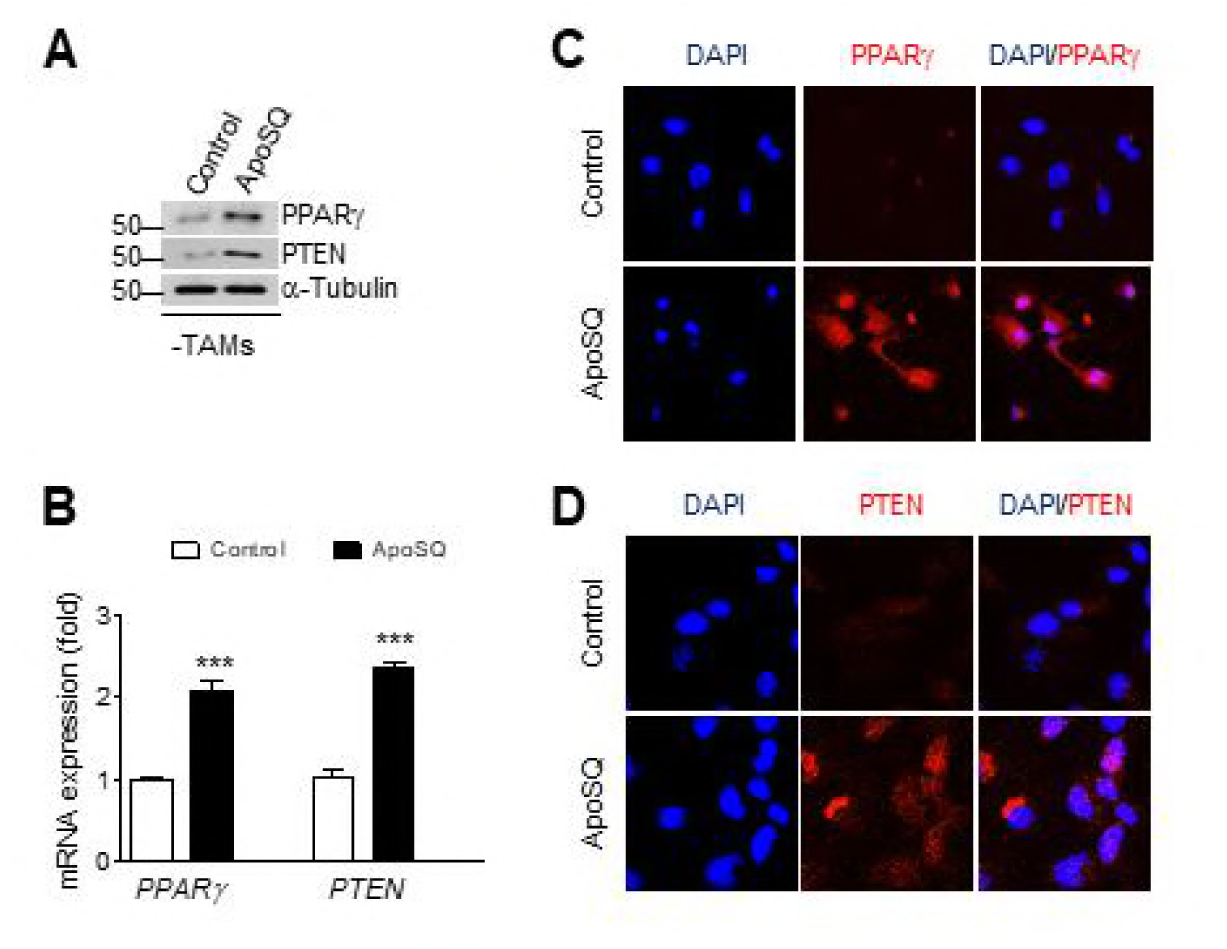
PPARγ and PTEN expression in tumor cells. Apoptotic 344SQ cells (ApoSQ) were subcutaneously injected in the skin lesion after subcutaneous injection of 344SQ cells into syngeneic (129/ν) mice (n=8 per group). Mice were necropsied 6 weeks later. (A) Immunoblot analysis and (B) qPCR analysis of remainder cells, lacking tumor-associated macrophages (TAM), isolated from primary tumors. (C and D) Representative confocal images of cells. \*\*\**P* < 0.001 (Student’s t test). Data represent mean ± s.e.m. from 5 mice.

**Supplementary figure 9.**
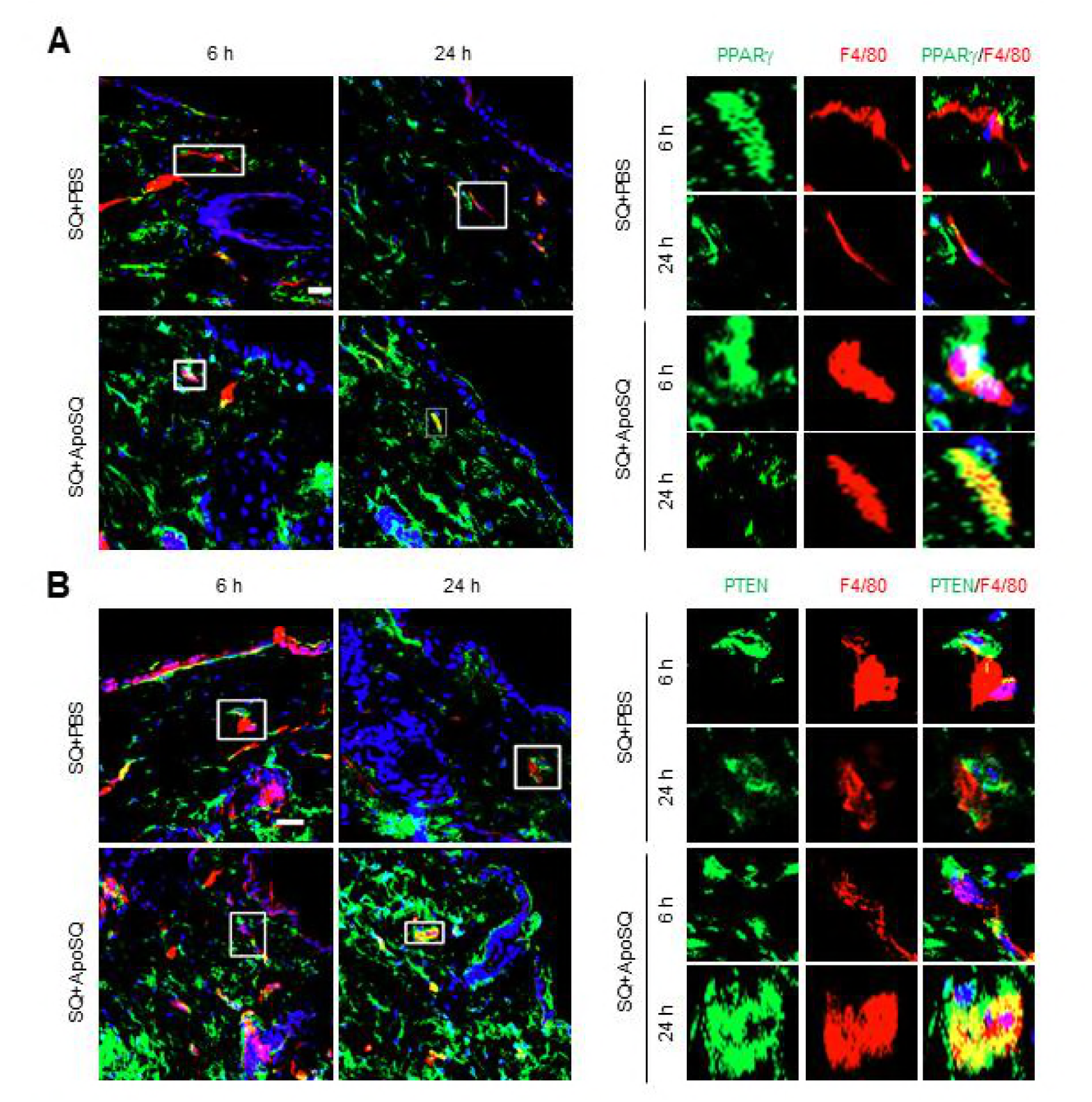
PPARγ and PTEN induction in subcutaneous macrophages following apoptotic 344SQ cell injection. Phosphate-buffered saline (PBS) or apoptotic 344SQ cells (ApoSQ) were subcutaneously injected in the skin lesion 2 days after subcutaneous injection of 344SQ cells into wild-type B6129SF2/J mice. (A and B) Mice were necropsied 6 or 24 h later. Confocal images of macrophages expressing PPARγ and PTEN in skin lesions immunostained with anti-F4/80 (red; A and B), anti-PPARγ (green; A), or anti-PTEN (green; B). Full-size images of skin lesions (A and B *left panels*) and enlarged ROIs from white squares on left panels (A and B *right panels*). Data are representative images from three mice per group.

**Supplementary table 1.** List of antibodies used this study.

| Antigen | Vendor | Cat. No | Source | Species cross-reactivity | Application | Dilution |
| --- | --- | --- | --- | --- | --- | --- |
| PPARγ | Santa Cruz Biotechnology | sc-7273 | Mouse monoclonal | H, M | IB, IF | 1:1000, 1:50 |
| ERK1/2 | Santa Cruz Biotechnology | sc-93-G | Goat polyclonal | H, M, R, C, F | IB | 1:1000 |
| P-ERK1/2 | Santa Cruz Biotechnology | sc-7383 | Mouse monoclonal | H, M | IB | 1:1000 |
| AKT | Santa Cruz Biotechnology | sc-8312 | Rabbit polyclonal | H, M, R, X | IB | 1:1000 |
| P-AKT | Santa Cruz Biotechnology | sc-7985-R | Rabbit polyclonal | H, M, R, X | IB | 1:1000 |
| p38 | Santa Cruz Biotechnology | sc-535-G | Goat polyclonal | H, M, R | IB | 1:1000 |
| P-p38 | Santa Cruz Biotechnology | sc-17852-R | Rabbit polyclonal | H, M, R | IB | 1:1000 |
| GFP | Santa Cruz Biotechnology | sc-9996 | Mouse monoclonal | Not Applicable | IB | 1:1000 |
| Na<sup>+</sup>/K<sup>+</sup> ATPase | Cell Signaling Technology | 3010 | Rabbit polyclonal | H, M, R, Mk, Hm, Z | IB | 1:1000 |
| PTEN | Cell Signaling Technology | 9559 | Rabbit monoclonal | H, M, R, Mk | IB, IP | 1:1000, 1:100 |
| E-cadherin | Cell Signaling Technology | 14472 | Mouse monoclonal | H, M, R | IB | 1:1000 |
| N-cadherin | Cell Signaling Technology | 4061 | Rabbit polyclonal | H, M, R, Mk | IB | 1:1000 |
| Vimentin | Cell Signaling Technology | 5741 | Rabbit monoclonal | H, M, R, Mk | IB | 1:1000 |
| P-Smad2 | Cell Signaling Technology | 3108 | Rabbit monoclonal | H, M, R, Mi | IB | 1:1000 |
| P-Smad3 | Cell Signaling Technology | 9520 | Rabbit monoclonal | H, M, R | IB | 1:1000 |
| α-Tubulin | Sigma-Aldrich | T5168 | Mouse monoclonal | H, M, R, Mk, C | IB | 1:2000 |
| PTEN | Abcam | ab170941 | Rabbit monoclonal | H, M, R | IF | 1:50 |
| CD36 | Abcam | ab80080 | Rat monoclonal | H, M | IF | 1:100 |
| CD63 | Abcam | ab193349 | Mouse monoclonal | H, M, Ct, Cw, Rb | IB | 1:1000 |
| F4/80 | Abcam | ab111101 | Rabbit monoclonal | M, R | IF | 1:100 |
| L-PGDS | Abcam | ab182141 | Rabbit monoclonal | H, M, R | IB | 1:1000 |
| β-catenin | BD Biosciences | 610154 | Mouse monoclonal | H, M, R, Dg, C | IF | 1:100 |
| Smad2/3 | BD Biosciences | 610842 | Mouse monoclonal | H, M, R, Dg | IB | 1:1000 |
| Mouse IgG (HRP) | Santa Cruz Biotechnology | sc-2005 | Goat | Not Applicable | IB | 1:5000 |
| Rabbit IgG (HRP) | Santa Cruz Biotechnology | sc-2004 | Goat | Not Applicable | IB | 1:5000 |
| Goat IgG (HRP) | Santa Cruz Biotechnology | sc-2354 | Mouse | Not Applicable | IB | 1:5000 |
| Rat IgG H&L<br>(Alexa Fluor647) | Abcam | ab150159 | Goat | Not Applicable | IF | 1:100 |
| Mouse IgG H&L<br>(Alexa Fluor 488) | Thermo Fisher Scientific | A11029 | Goat | Not Applicable | IF | 1:100 |
| Rabbit IgG H&L<br>(Alexa Fluor 488) | Thermo Fisher Scientific | A11034 | Goat | Not Applicable | IF | 1:100 |
| Mouse IgG H&L<br>(Alexa Fluor 594) | Thermo Fisher Scientific | A11005 | Goat | Not Applicable | IF | 1:100 |
| Rabbit IgG H&L<br>(Alexa Fluor 594) | Thermo Fisher Scientific | A11012 | Goat | Not Applicable | IF | 1:100 |

Abbreviation: IB-Immunoblot, IP-Immunoprecipitation, IF-Immunofluorescence, H-human, M-mouse, R-rat, Hm-hamster, Mk-monkey, Mi-mink, C-chicken, Ct-cat, Cw-cow, Dg-dog, F-frog, Rb-rabbit, X-Xenopus, Z-zebrafish

**Supplementary table 2.**
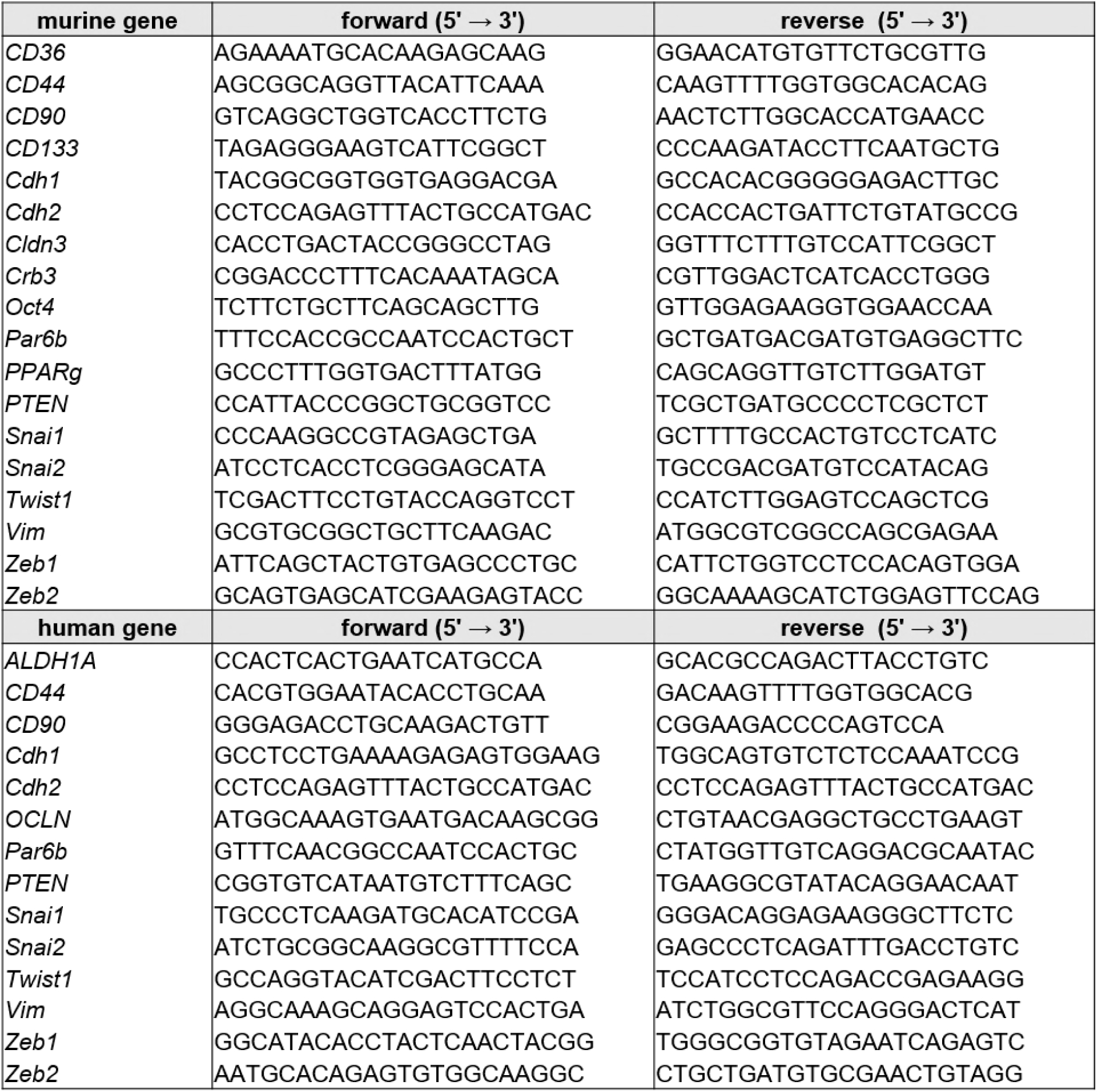
qPCR primers used in this study.

